# VAE-SNE: a deep generative model for simultaneous dimensionality reduction and clustering

**DOI:** 10.1101/2020.07.17.207993

**Authors:** Jacob M. Graving, Iain D. Couzin

## Abstract

Scientific datasets are growing rapidly in scale and complexity. Consequently, the task of understanding these data to answer scientific questions increasingly requires the use of compression algorithms that reduce dimensionality by combining correlated features and cluster similar observations to summarize large datasets. Here we introduce a method for both dimension reduction and clustering called VAE-SNE (variational autoencoder stochastic neighbor embedding). Our model combines elements from deep learning, probabilistic inference, and manifold learning to produce interpretable compressed representations while also readily scaling to tens-of-millions of observations. Unlike existing methods, VAE-SNE simultaneously compresses high-dimensional data and automatically learns a distribution of clusters within the data — without the need to manually select the number of clusters. This naturally creates a multi-scale representation, which makes it straightforward to generate coarse-grained descriptions for large subsets of related observations and select specific regions of interest for further analysis. VAE-SNE can also quickly and easily embed new samples, detect outliers, and can be optimized with small batches of data, which makes it possible to compress datasets that are otherwise too large to fit into memory. We evaluate VAE-SNE as a general purpose method for dimensionality reduction by applying it to multiple real-world datasets and by comparing its performance with existing methods for dimensionality reduction. We find that VAE-SNE produces high-quality compressed representations with results that are on par with existing nonlinear dimensionality reduction algorithms. As a practical example, we demonstrate how the cluster distribution learned by VAE-SNE can be used for unsupervised action recognition to detect and classify repeated motifs of stereotyped behavior in high-dimensional timeseries data. Finally, we also introduce variants of VAE-SNE for embedding data in polar (spherical) coordinates and for embedding image data from raw pixels. VAE-SNE is a robust, feature-rich, and scalable method with broad applicability to a range of datasets in the life sciences and beyond.

## 1 Introduction

Modern scientific research generates large, high-resolution datasets that are complex and high-dimensional, where a single observation from an experimental system can contain measurements describing hundreds, or thousands, of features. For example, neuroscientists measure electrical activity across thousands of individual neurons simultaneously (Jun et al., 2017; Stringer et al., 2019a,b) — even across the entire brain (Ahrens et al., 2012, 2013); cell biologists and bioinformaticians routinely sequence the transcriptome for thousands of genes across large populations of single cells (Samusik et al., 2016; La Manno et al., 2018; Becht et al., 2019; Linderman et al., 2019); behavioral scientists measure the high-dimensional body posture dynamics of animals and humans (Stephens et al., 2008, 2011; Kain et al., 2013; Berman et al., 2014; Wiltschko et al., 2015; Klibaite et al., 2017; Costa et al., 2019; Cande et al., 2018; Mathis et al., 2018; Chambers et al., 2019; Günel et al., 2019; Graving et al., 2019; Klibaite and Shaevitz, 2019; Nath et al., 2019; Pereira et al., 2019; Bala et al., 2020; Ebbesen and Froemke, 2020; Karashchuk et al., 2020); and evolutionary ecologists measure complex morphological patterns across sizeable collections of animal specimens (Cuthill et al., 2017, 2019; Ezray et al., 2019; Wham et al., 2019; Zhang et al., 2019). While there are many benefits to measuring real-world systems accurately and completely for answering scientific questions, this added complexity poses problems for conventional data analysis methods — especially those commonly used in the life sciences, like linear models (Bolker et al., 2009) — that are designed for small, low-dimensional datasets and typically rely on simplified models with strong, often unrealistic, assumptions for making statistical inferences.

To deal with the complexity of modern data, researchers in many fields have begun to use machine-learning methods known as *dimensionality reduction* and *clustering* to help interpret large, high-dimensional datasets. These algorithms distill correlated features down to a smaller set of components (dimensionality reduction) or group large subsets of observations into a smaller set of classes based on similarity (clustering). Together these methods offer scientists a way to *compress* data, where compression is typically performed with the goal of reducing the size and complexity of a dataset while making only minimal, or very general, a priori assumptions about the true distribution of the data. Because these algorithms derive their compressed representations directly from the structure of the data itself, without human supervision, they are typically known as *unsupervised learning* algorithms.

Across many scientific disciplines, unsupervised algorithms are rapidly becoming a commonly-used tool for visualizing and interpreting high-dimensional data distributions as well as summarizing large datasets with coarse-grained descriptions and identifying specific subpopulations and regions of interest within the data for further downstream analysis. Researchers have applied these methods to demonstrate how the brain organizes behavior (Stephens et al., 2008, 2011; Brown et al., 2013; Wiltschko et al., 2015; Berman et al., 2016; Billings et al., 2017; Cande et al., 2018; Markowitz et al., 2018; Costa et al., 2019; Stringer et al., 2019a, b); describe how cells grow and develop over time (La Manno et al., 2018); document new and rare types of cells (Grün et al., 2015; Linderman et al., 2019); gain insights into cancer treatment (Tirosh et al., 2016); and reveal fundamental principles of evolution (Cuthill et al., 2019; Ezray et al., 2019; Wham et al., 2019). Therefore, as scientists begin to regularly rely on these algorithms for analyzing complex datasets, the task of ensuring the quality, robustness, and utility of the compressed representations they produce is an issue of considerable importance — as is the ability to scale these methods to increasingly large datasets.

While existing methods for dimension reduction produce high-quality compressed representations (Becht et al., 2019; Kobak and Linderman, 2019), they typically lack features for identifying groups of similar data (i.e., learned clusters; but see Pezzotti et al.2016; Robinson and Pierce-Hoffman 2020), and despite much progress to improve scalability of existing algorithms (Linderman et al., 2017; McInnes et al., 2018; Linderman et al., 2019), some of the most widely-used methods are still limited in their ability to scale beyond a few million observations without specialized, high-performance hardware — especially in the case of large, out-of-core datasets that cannot fit into memory. Recent applications of deep learning (Goodfellow et al., 2016), and deep generative models in particular (Appendix A.1; Kingma and Welling 2013; Rezende et al.2014), have begun to address these issues (Ding et al., 2018; Szubert et al., 2019; Ding and Regev, 2019). Nevertheless, even with the low memory and computational cost of deep learning methods that can be trained with small batches of data on consumer-grade hardware, these new algorithms are still significantly slower to fit to data than more popular methods because they require costly nearest neighbor or pairwise distance calculations (Becht et al., 2019; Ding et al., 2018; Szubert et al., 2019). The majority of these methods also do not provide any built-in mechanism for detecting outliers, which could potentially bias any downstream results and cause statistical errors when testing hypotheses.

There has also been a flurry of recent work on advanced methods for clustering data (e.g., Campello et al. 2013; Jiang et al. 2016; Xie et al. 2016; Guo et al. 2017; McInnes et al. 2017; Fogel et al. 2019; Yang et al. 2019; Robinson and Pierce-Hoffman 2020; and numerous others), including efficient methods that rely on deep learning and deep generative models. However, the vast majority of these methods impose strong assumptions about the shape of the clusters and require the user to manually select the number of clusters fitted to the data — or, alternatively, involve complex computations that do not scale well to large datasets. Determining how many clusters to fit is typically a non-trivial, unintuitive, and computationally-intensive task for datasets where the number of clusters is not known a priori (Milligan and Cooper, 1985; Pham et al., 2005; Fang and Wang, 2012; Todd et al., 2017). Many recently proposed clustering algorithms are also only evaluated with relatively small “toy” datasets, such as the MNIST handwritten digit database (LeCun et al., 2010), where the data typically have very little noise, no outliers, and the number of clusters is often known a priori. This lack of rigorous real-world assessment casts doubt on the practical utility of these algorithms in cases where datasets have a large number of observations, are naturally noisy or contain outliers, and the number of clusters is unknown, such as those commonly used in the natural sciences.

Here we aim to address many of the limitations outlined above and unify some of the key methodological concepts from previous work into a single modeling framework. To accomplish this, we introduce a deep generative model for both dimensionality reduction and clustering. We then compare our model with existing methods for dimensionality reduction, and importantly, to ensure that it has practical utility, we demonstrate the application of our method using empirical examples with real-world data from multiple domains. In comparison to existing dimension reduction methods, our proposed method produces low-dimensional data representations with similar, or better, quality while also offering several key improvements. Notably, our approach provides the ability to scale to datasets containing tens-of-millions of observations without specialized, high-performance hardware and automatically learns an interpretable cluster distribution from the data without any manual tuning or expensive computations to determine the number of clusters. Together these results demonstrate that our proposed method is a robust, feature-rich, and scalable tool for data analysis and is widely-applicable to a variety of tasks.

## 2 Results

We make three main contributions in this paper: (**1**) First, we introduce a deep generative model for both dimensionality reduction and clustering called variational autoencoder stochastic neighbor embedding (VAE-SNE; Fig. 1; Methods). VAE-SNE can produce a variety of different compressed representations and readily scales to out-of-core datasets with tens-of-millions of observations. Our model builds on numerous ideas from past work by synthesizing methods from a class of generative models known as variational autoencoders (VAEs; Kingma and Welling 2013), the popular dimensionality reduction algorithm (*t*-distributed) stochastic neighbor embedding (SNE/t-SNE; Hinton and Roweis 2003; van der Maaten and Hinton 2008) and its many extensions (van der Maaten, 2009; Wang and Wang, 2016; Chien and Hsu, 2017; Ding et al., 2018), as well as recent advances in variational inference (Kingma et al., 2014; Burda et al., 2015; Dilokthanakul et al., 2016; Cremer et al., 2017; Tomczak and Welling, 2017) and clustering methods (Todd et al., 2017). (**2**) Second, we apply VAE-SNE, and a variety of other popular dimensionality reduction methods, to compress real-world datasets from different domains (Fig. 2). We then quantitatively assess how each algorithm performs in preserving important aspects of the data — including information about local, global, and temporal structure. We also assess generalization to new, out-of-sample data and compare processing speeds for each algorithm. Additionally, we show how the likelihood score produced by VAE-SNE can be used to detect outliers when embedding out-of-sample data. (**3**) Third, we show how VAE-SNE can be used to automatically cluster large datasets into a small set of interpretable classes. As a practical example, we apply VAE-SNE to a dataset of 21.1 million observations describing the high-dimensional body posture dynamics of a commonly-used model organism — the fruit fly (*Drosophila melanogaster*) — to automatically discretize these data into motifs of stereotyped behavior for further analysis (Fig. 3; Berman et al. 2014; Pereira et al. 2019). These results illustrate how VAE-SNE can be used as a type of automated ethogram for describing the full behavioral repertoire of animals (reviewed by Anderson and Perona 2014; Berman 2018; Brown and De Bivort 2018; Datta et al. 2019), while also providing several advantages over existing methods for this task.

**Figure 1.**
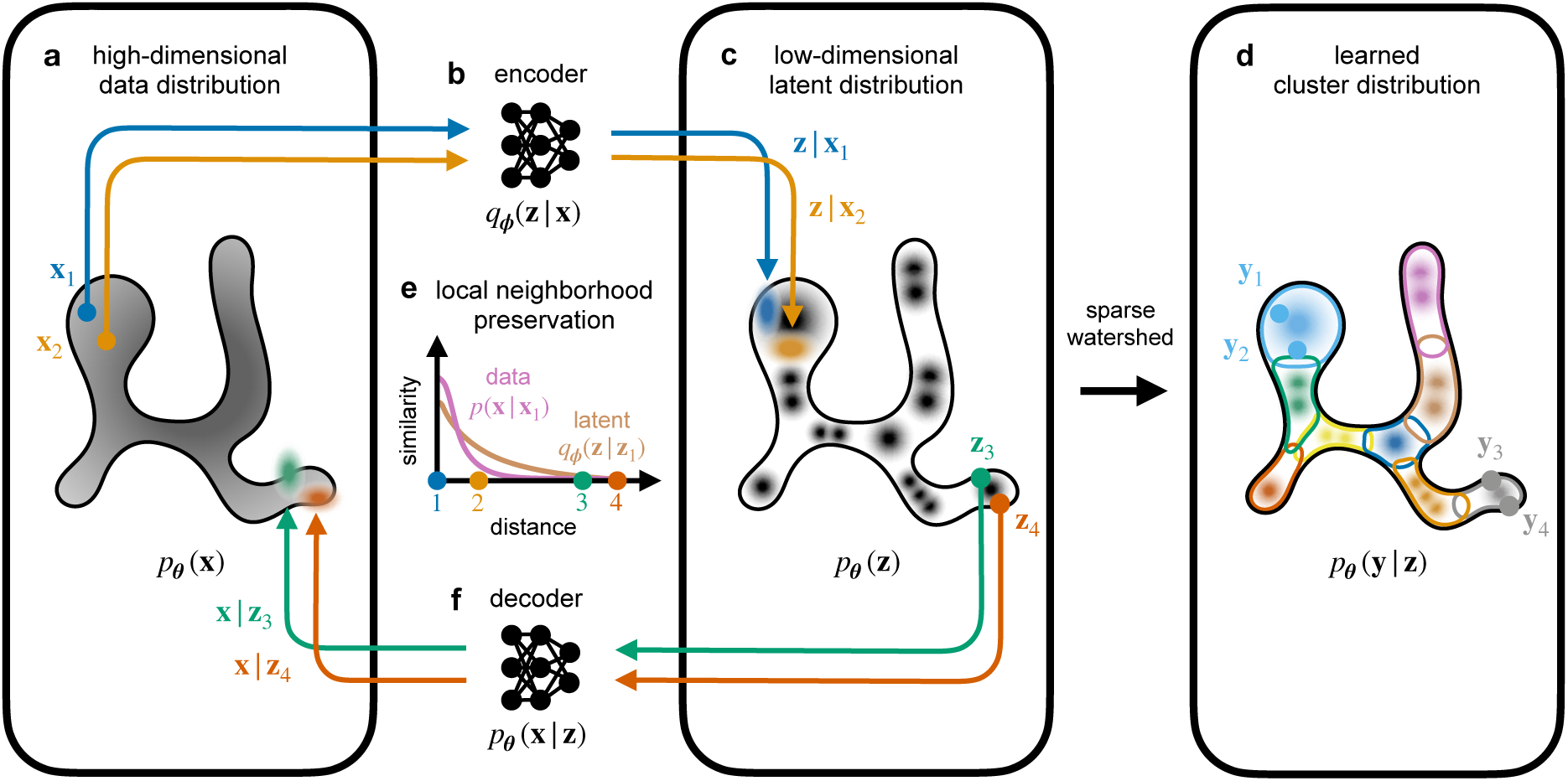
Overview of the VAE-SNE model. **a**-**f**, Observed samples from a high-dimensional data distribution **x** ∼ *p*(**x**) (**a**) are probabilistically embedded (**b**) into a low-dimensional latent distribution *p*_*θ*_(**z**) (**c**) using an encoder deep neural network DNN_*ϕ*_ : **x** → **z** to generate an approximate latent posterior distribution *q*_*ϕ*_(**z**|**x**). Samples from the latent distribution **z** ∼ *q*_*ϕ*_(**z**|**x**) or **z** ∼ *p*_*θ*_(**z**) (**c**) are then transformed (**f**) using a generative decoder deep neural network DNN_*θ*_ : **z** → **x** to probabilistically reconstruct the high-dimensional data distribution *p*_*θ*_(**x**|**z**). Given a set of observed high-dimensional data {**x**_1_, **x**_2_, …, **x**_*N*_} the model parameters for the encoder and decoder {***θ***, *ϕ*} are optimized so that the approximate posterior for the encoder matches the true posterior from the generative decoder as best as possible, or *q*_*ϕ*_(**z**|**x**) ≈ *p*_*θ*_(**z**|**x**), which then creates a functional mapping between the high-dimensional and low-dimensional distributions. To improve local structure preservation during optimization, pairwise distances between vectors in the high-dimensional and low-dimensional space are optimized using pairwise similarity kernels (**e**), a probability density function of distance, so that the local neighborhoods around each observation match as best as possible, or *p*(**x**|**x**_*i*_) ≈ *q*_*ϕ*_(**z**|**z**_*i*_). This preferentially weights the preservation of local neighborhoods over global relationships by assigning more probability mass to nearby neighbors during optimization. The prior for the latent distribution *p*_*θ*_(**z**) is also a learned Gaussian mixture distribution (**c**) that is jointly optimized with the encoder and decoder to fit the observed data and can be used to cluster the latent distribution (**d**) into a small set of discrete classes *p*_*θ*_(**y**|**z**) — where highly-overlapping modes (mixture components) within the distribution are automatically merged into the same class label using sparse watershed assignment (Methods; Todd et al. 2017)

**Figure 2.**
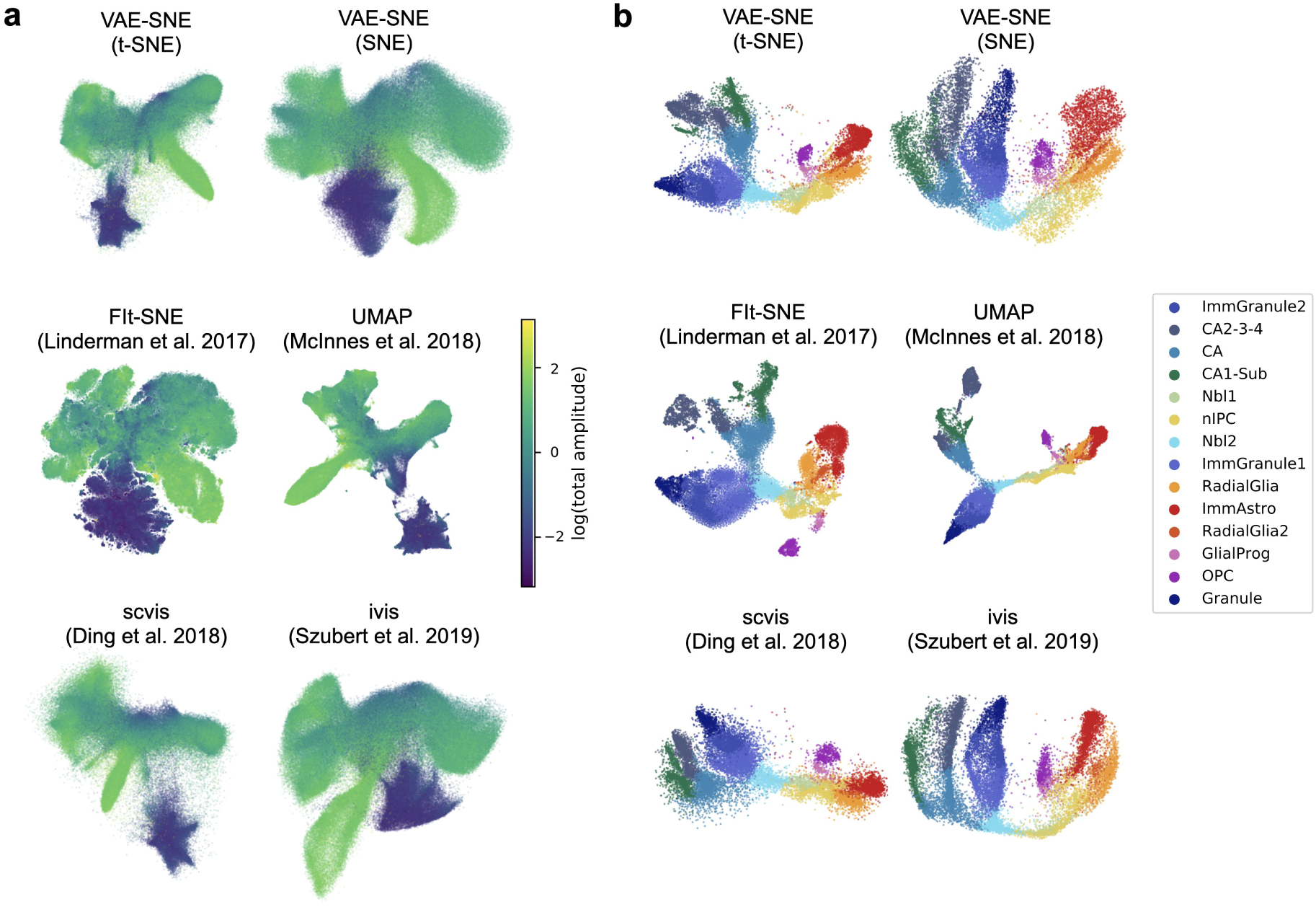
Embeddings for body posture dynamics and single-cell RNA-seq data. **a**, 2-D embeddings of body posture dynamics data from Berman et al. (2014, 2016); Pereira et al. (2019) for each algorithm we tested. The color of each point indicates the logarithm of the total amplitude (overall movement) of body parts for each observation. **b**, 2-D embeddings of single-cell RNA-seq data of developing hippocampal neurons from La Manno et al. (2018) for each algorithm. The color of each point indicates the cell type for that observation as described by La Manno et al. (2018).

**Figure 3.**
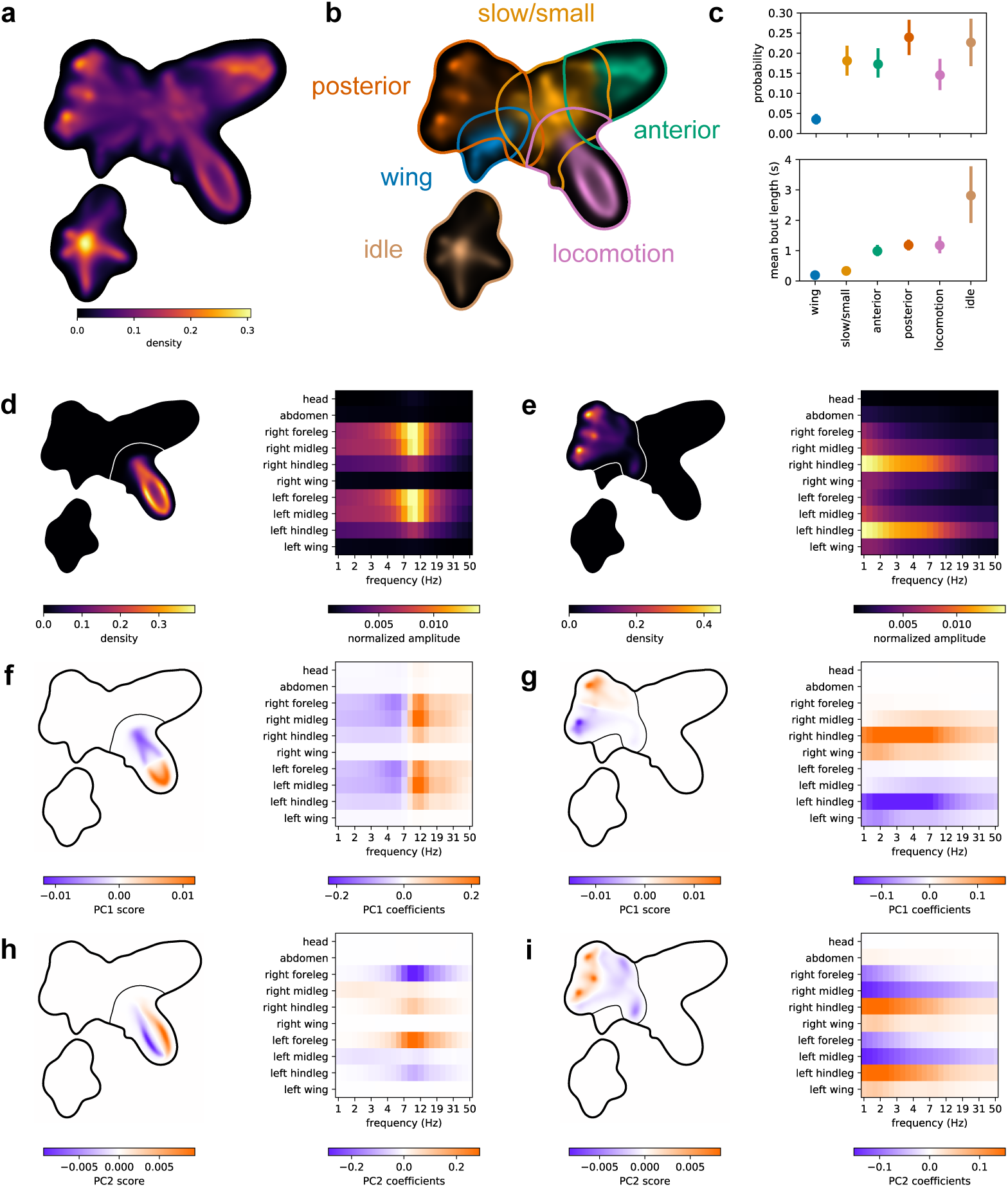
Clustering body posture dynamics. **a**, The posterior probability density for the full 21.1 million observation body posture dynamics dataset from Berman et al. (2014, 2016); Pereira et al. (2019) embedded using a 2-dimensional VAE-SNE model. **b**, The manually-grouped high-level cluster assignments produced using the learned prior from a 30-dimensional VAE-SNE embedding visualized in the 2-D embedding, where contours are the largest 90% probability density contour for each cluster distribution. **c**, Mean and 95% bootstrap intervals of the marginal (stationary) probability and mean bout length for each high-level cluster (n = 59 per cluster). **d**-**i**, Visualizations describing the high-level locomotion (**d**,**f**,**h**; Video S2; Fig. S8) and posterior grooming (**e**,**g**,**i**; Video S4; Fig. S9) clusters. **d-e**, The 2-D posterior probability density for each cluster (left) and the mean spectrogram for each cluster (right). **f**-**i**, The principal component scores for the two largest components of the spectrograms assigned to each cluster visualized within the 2-D embedding (left), and the eigenvector coefficients describing the linear contribution of each spectrogram feature (right) for the principal component score.

Our approach (Fig. 1; Methods) builds on VAEs as a base model for performing dimensionality reduction (Appendix A.1), which, like other types of autoencoders (Hinton and Salakhutdinov, 2006), model high-dimensional data using two deep neural networks: one to encode data to a compressed latent representation, and another to decode the latent vectors and reconstruct the data. However, VAEs are distinct from other autoencoders in that the encoder is used to parameterize continuous distributions of latent vectors — from which latent vectors are then probabilistically sampled — rather than embedding each high-dimensional observation as a single point in the latent space. This type of model offers an attractive dimensionality reduction framework because the objective function (Appendix A.2) naturally imparts a trade-off between the complexity of the encoded description and the overall accuracy of the decoded reconstruction (Alemi et al., 2016). However, these models suffer from multiple long-standing issues including a phenomenon known as *posterior collapse* (Alemi et al., 2017; Dieng et al., 2019a) where the latent coordinate space becomes arbitrarily organized and no longer preserves any statistical features of the high-dimensional data distribution. There has been a string of recent work to address these issues including some relatively straightforward solutions (Higgins et al., 2016; Dieng et al., 2019a) that achieve varying levels of success, as well as new objective functions that involve regularizing the mutual information between the high-dimensional data and latent distribution (e.g., Zhao et al. 2017; Rezaabad and Vishwanath 2019; reviewed by Poole et al. 2019).

For VAE-SNE, we provide an effective solution to this problem with the addition of a stochastic neighbor regularizer (Appendix B; van der Maaten and Hinton 2008; van der Maaten 2009; Chien and Hsu 2017; Ding et al. 2018) that optimizes pairwise similarity kernels between the high- and low-dimensional distributions to strengthen local neighborhood preservation and more explicitly retain a useful representation. We also draw on other theoretical and practical improvements from the literature to enhance the performance of VAE-SNE (Methods). For example, we use a Gaussian mixture prior for learning the latent distribution (Kingma et al., 2014; Dilokthanakul et al., 2016; Tomczak and Welling, 2017). This choice of distribution allows for better local structure preservation and, when combined with sparse watershed assignment to merge overlapping mixture components (Fig. 1; Methods; Todd et al. 2017), serves as a flexible method for clustering data — without the need to manually define the number of clusters or impose strong assumptions about cluster shape. We employ several other advances to further improve structure preservation. For instance, we apply a perplexity annealing technique (Kobak and Berens, 2019) to slowly decay the size of the local neighborhoods optimized by the model during training, which helps to preserve structure across multiple scales. Moreover, we extensively optimize the algorithms underlying our model by applying parallel computations on the CPU and GPU that dramatically improve processing speed compared to previous work (Ding et al., 2018).

In addition to our three main contributions, we further extend VAE-SNE to demonstrate its flexibility as a framework for dimensionality reduction. To accomplish this, we introduce a von Mises-Fisher variant of VAE-SNE (Appendix C.1; Fig. S10; Video S8,Video S9) that embeds data in polar coordinates (rather than Euclidean coordinates) on a 3-D unit sphere, which is potentially a more natural representation for many high-dimensional datasets (Davidson et al., 2018) and solves the “crowding” problem common to some methods (van der Maaten and Hinton, 2008; Ding and Regev, 2019). Finally, we also apply a modified convolutional version of VAE-SNE (Appendix C.2; Figs. S11, S12) to visualize natural history images of animal specimen collections (Cuthill et al., 2019; Zhang et al., 2019) by directly embedding the raw pixel data. Our results for these two extensions are described in Appendix C.

### 2.1 Comparisons with other dimension reduction algorithms

Current methods for dimensionality reduction generally fall into two classes known as *linear* and *nonlinear* algorithms. Linear algorithms, such as principal components analysis (PCA), compress high-dimensional data by learning linearly weighted combinations (affine transformations) of the original feature set. Typically these algorithms are optimized to preserve the global structure of the data, where local neighborhood relationships are distorted in order to maintain the full coordinate system of the original features as best as possible. On the other hand, nonlinear algorithms (sometimes called manifold learning algorithms) such as t-SNE (van der Maaten and Hinton 2008) and uniform manifold approximation and projection (UMAP; McInnes et al. 2018) typically take the opposite approach of prioritizing relative relationships between data points rather than the global coordinate system. This approach allows local neighborhoods to be preserved while potentially sacrificing information about the larger-scale relationships between data points in the global coordinate space — although, as we demonstrate here, the global distortion imposed by many of these algorithms is actually comparable to that of PCA.

To validate VAE-SNE as a general-purpose method for dimensionality reduction, we quantitatively compare its performance with other dimension reduction algorithms — both linear and nonlinear — using two datasets from different domains (see Methods) describing animal body part dynamics (Berman et al., 2014, 2016; Pereira et al., 2019) and single-cell RNA-seq expression profiles for hippocampal neurons (La Manno et al., 2018). We benchmark multiple variants of VAE-SNE with different pairwise similarity kernels for preserving local neighborhood information (including kernel functions with learned parameters; Appendix B), and we compare these results with those from two high-performance variants of t-SNE (van der Maaten and Hinton, 2008) known as FIt-SNE (Linderman et al., 2017, 2019) and Barnes-Hut-SNE (van der Maaten, 2014), as well as UMAP (McInnes et al., 2018), and two other deep neural network-based dimension reduction methods: scvis (Ding et al., 2018), and ivis (Szubert et al., 2019). We also apply PCA in 2, 5, 10, and 100 dimensions for a linear baseline comparison. We fit each algorithm with a training set and also embed an out-of-sample test set to assess generalization to new data. For both the training and test sets, we then quantitatively assess each algorithm’s ability to preserve different types of information about the high-dimensional data when compressing the data to two dimensions, including local, global, fine-scale, and temporal information (Methods). We quantify local information preservation for each algorithm by measuring the preservation of both metric (distance- or radius-based) and topological (nearest neighbors-based) neighborhoods that are approximately 1% of the total embedding size; we measure global information preservation by calculating the correlation between pairwise distances in high- and low-dimensional space; we assess fine-scale information by measuring neighborhood preservation for multiple neighborhood sizes < 1% of the total embedding size; and we evaluate temporal information preservation by computing the correlation between high- and low-dimensional temporal derivatives in a timeseries dataset. Overall the qualitative properties of the embeddings produced by each algorithm are strikingly similar within datasets (Fig. 2), which likely indicates shared mathematical properties of how the latent distributions are modelled. However, we do find potentially important quantitative differences between these algorithms in terms of information preservation and processing speed. We summarize our overall assessments of each nonlinear dimension reduction algorithm in Tables S1, S2, S3.

#### 2.1.1 Local structure preservation

We find that VAE-SNE compares closely to FIt-SNE (Linderman et al., 2017), Barnes-Hut-SNE (van der Maaten, 2014), and UMAP (McInnes et al., 2018) in preserving local structure for both the training set (Figs. S1a, S2a, S5a) and test set (Figs. S3a, S4a), while scvis (Ding et al., 2018) and ivis (Szubert et al., 2019) perform slightly worse. Our results show that VAE-SNE with a t-SNE similarity kernel (van der Maaten and Hinton, 2008) performs the best for preserving local structure, but VAE-SNE with a Gaussian SNE kernel (Hinton and Roweis, 2003) also performs well — similarly to scvis (Ding et al., 2018) and ivis (Szubert et al., 2019). We also find that learning the similarity kernel parameters (for both Gaussian and Student’s *t* kernels) as a function of each data point does not improve performance for our local preservation metrics. The top performing algorithms for local structure preservation (VAE-SNE, t-SNE, and UMAP) are closely comparable to 5-dimensional PCA for both metrics we used to assess local neighborhood preservation.

#### 2.1.2 Global structure preservation

We find that VAE-SNE also does well in preserving global structure for both the training set (Figs. S1a, S2b, S5a) and test set (Figs. S3a, S4b). VAE-SNE with a Gaussian SNE kernel performs best for this metric, but VAE-SNE with a t-SNE kernel also performs nearly as well. Notably all the neural-network-based methods (VAE-SNE, scvis Ding et al. 2018, ivis Szubert et al. 2019) outperform both t-SNE and UMAP (McInnes et al., 2018) in preserving global structure for both datasets we tested. This is perhaps not surprising given that recent work has shown neural network models tend to learn the same axes as PCA (Rolinek et al., 2019). Additionally, these results show that learning the similarity kernel parameters as a function of each data point does improve global structure preservation for VAE-SNE with a t-SNE kernel — likely because it is optimized to be more similar to the Gaussian kernel used to calculate high-dimensional similarities (Appendix B). The top performing algorithms for this metric are comparable to 2-dimensional PCA, which demonstrates that nonlinear algorithms are capable of preserving the same global information as PCA while also better preserving local structure. On one hand, The scvis (Ding et al., 2018) algorithm in particular excels at preserving global structure for the single-cell RNA-seq dataset we tested (Fig. S5a). On the other hand, ivis (Szubert et al., 2019) performs much more poorly than the other neural network algorithms for this dataset, and FIt-SNE (Linderman et al., 2017, 2019) and Barnes-Hut-SNE (van der Maaten, 2014) perform even worse. We also show that UMAP (McInnes et al., 2018) with PCA initialization better preserves global structure than the default Laplacian Eigenmap initialization.

#### 2.1.3 Fine-scale structure preservation

In addition to local and global structure preservation, we evaluate the ability of each algorithm to preserve very fine-scale neighborhood information (Figs. S1b, S3b, S5b). We find that both FIt-SNE (Linderman et al., 2017) and Barnes-Hut-SNE (van der Maaten, 2014) excel at preserving this fine-scale information for the posture dynamics dataset (Figs. S1b, S3b) while every other nonlinear algorithm performs relatively poorly for both the training and test set. For the single-cell RNA-seq dataset, this distinction is not nearly as large and the algorithms all perform more similarly (Fig. S5b), which indicates performance varies depending on the dataset. Performance for the ivis algorithm (Szubert et al., 2019) is especially poor for this metric on the single cell RNA-seq dataset. However, neighborhood membership for neighborhoods between 1% and 10% of the total embedding size are all similarly well-preserved for each algorithm.

#### 2.1.4 Temporal structure preservation

Because one of the datasets we use for benchmarking is a behavioral timeseries, for these data we also assess the temporal structure preservation of each algorithm (Figs. S3a, S4c) on the out-of-sample test set (the training set is randomly sampled across multiple timeseries, so temporal information is not preserved). We find that VAE-SNE (particularly the SNE kernel variant), FIt-SNE (Linderman et al., 2017), Barnes-Hut-SNE (van der Maaten, 2014), scvis (Ding et al., 2018), and ivis (Szubert et al., 2019) perform at the same level as 5-dimensional PCA in preserving temporal structure, while UMAP (McInnes et al., 2018) performs relatively poorly in comparison to the other algorithms — even worse than 2-dimensional PCA.

#### 2.1.5 Speed comparisons

In addition to assessing information preservation, we also compare the speed the of each algorithm both when fitting the algorithm to the training set (Figs. S1c, S5c) and when embedding an out-of-sample test set (Figs. S3c, S5c). We find that training time increases approximately linearly with the size of the dataset for each algorithm. UMAP (McInnes et al., 2018) has the fastest training time (approximately as fast as PCA), followed by FIt-SNE (Linderman et al., 2017) and Barnes-Hut-SNE (van der Maaten, 2014), and then VAE-SNE. While VAE-SNE is slower for fitting the training set than both UMAP (McInnes et al., 2018) and t-SNE, it is much faster than the other two neural network methods scvis (Ding et al., 2018) and ivis (Szubert et al., 2019). We also demonstrate that VAE-SNE, and the other neural network methods, can quickly embed out-of-sample test data (Figs. S3c, S5c). The time needed for embedding new data is much higher for both t-SNE and UMAP, and while the elapsed time for embedding the test set scales linearly with the number of samples for all algorithms, we also find that it increases with the size of the training set for both UMAP (McInnes et al., 2018) and Barnes-Hut-SNE (van der Maaten, 2014) (Fig. S3c). This is almost certainly because adding new data for these algorithms requires calculating approximate nearest neighbors between the out-of-sample data and the training set, which consequently requires more computation time for larger training sets. Unexpectedly, FIt-SNE (Linderman et al., 2017) does not exhibit this behavior despite using similar nearest neighbor calculations to Barnes-Hut-SNE (van der Maaten, 2014). On the other hand, VAE-SNE and other deep learning algorithms do not suffer from this limitation. Finally, while we do not comprehensively assess memory complexity of different algorithms in this paper, we stopped our speed comparisons at data subsets with 232,000 (× 1500 dimensions) observations because UMAP began to cause out-of-memory errors for larger subsets — while all of the other algorithms we tested could still successfully run under the same conditions. This helps to illustrate the key advantage of deep learning-based methods, which naturally maintain very low memory complexity by applying optimization using small batches of data.

### 2.2 Using the likelihood to assess out-of-sample data

Because VAE-SNE also calculates a likelihood score for reconstructing the original high-dimensional data, we can use this to assess performance on out-of-sample data, which is an idea originally proposed by Ding et al. (2018). To test this, we calculate the likelihood score for real data from the posture dynamics dataset (Berman et al., 2014, 2016; Pereira et al., 2019) and randomly-permuted data (randomized across feature columns) from the same dataset. We find that the likelihood score is reliably lower for the randomized data, and the two likelihood distributions are well separated (Fig. S6a), which shows this metric could potentially be used to detect outliers. We also compare the entropy of the approximate posterior distribution for each embedded sample as another potential metric for detecting outliers. While we find that the entropy is much higher for the randomized data, the distribution is highly overlapping with the entropy for the real data (Fig. S6b), which indicates the entropy may not be as useful for evaluating the embedding quality.

### 2.3 Clustering body posture dynamics to reveal stereotyped behavioral organization

To demonstrate its capabilities as a clustering algorithm, we use VAE-SNE to automatically discretize a dynamical time series dataset describing the high-dimensional body posture and behavioral repertoire of 59 freely-behaving fruit flies (*D. melanogaster*; Berman et al. 2014, 2016; Pereira et al. 2019) — a commonly-used model organism for neuroscience, pharmaceutical, and genetics research. To accomplish this, we use the annotated training data from (Pereira et al., 2019) to train a pose estimation model using deep learning-based software (DeepPoseKit; Graving et al. 2019). We then use this trained model to automatically track the spatial locations of 10 body parts (head, legs, wings, abdomen) directly from video timeseries data and generate time-frequency spectrograms describing body-part dynamics for each observation in the timeseries (Berman et al., 2014), which naturally incorporates multi-scale temporal information into each data vector. We then apply VAE-SNE to compress the data to a 30-dimensional latent embedding and simultaneously discretize the dynamical posture timeseries into a set of behavioral clusters. We find that, after optimizing the 30-D VAE-SNE model for 5 repeated trials using the full 21.1 million observation dataset and applying sparse watershed assignment to generate cluster labels (Methods; Fig. 1d; Todd et al. 2017), VAE-SNE consistently learns a total of 26 low-level behavioral clusters describing distinct, stereotyped body part movements. We also achieve similar (nearly identical) results when clustering in 10-D and 50-D space and when varying the number of components in the Gaussian mixture prior used for clustering — provided that the number of components is large enough (e.g., *K* ≥ 100).

To provide a broad overview of the behavioral structure discovered by VAE-SNE, we manually group these low-level clusters into 6 high-level clusters (Figs. 3, S7; Video S1) by examining video clips sampled from each cluster (Video S2–Video S7) and by calculating and visualizing the mean spectrograms for each low-level cluster to quantify the average distribution of body part movements across frequencies for each behavioral class (Figs. S8d-f, S9d-i). These high-level clusters include: locomotion (Video S2), anterior grooming (Video S3), posterior grooming (Video S4), wing movements (Video S5), small/slow leg movements (Video S6), and idle behavior (Video S7). Many of the low-level clusters (10 clusters in total) describe distinct slow/small leg movements, while there are 3 low-level clusters for locomotion (Fig. S8), 3 for anterior grooming, 6 for posterior grooming (Fig. S9), 2 for wing movements, and 2 for idle behavior. Videos and posture timeseries data sampled from each cluster also clearly demonstrate the stereotypy of behaviors within these behavioral classes, which matches well with previous work describing these dynamics (Berman et al., 2014, 2016; Klibaite et al., 2017; Klibaite and Shaevitz, 2019; Pereira et al., 2019). Additionally, the principal components of the spectrograms from each high-level cluster (Fig. 3f-i; Fig. S7d-i) reveal continuous variation related to asymmetrical body movements and differences in peak movement frequency. We calculate basic statistics describing cluster usage across individuals (Figs. 3c, S8c, S9c) including the marginal (stationary) probability of behavioral classes across individuals and the mean bout length, or the average amount of time a behavior is performed when an individual transitions into that cluster. In particular, the low probability and short bout length for wing movements and short bout length for slow/small leg movements (Fig. 3c) indicate these clusters may be transitional or idiosyncratic behaviors (Todd et al., 2017). For the low-level locomotion clusters (Fig. S8) we also calculate the forward component of the leg movement velocity (in body lengths per second, or BL · s^−1^) relative to the egocentric orientation of the animal. We then use the forward velocity to classify each leg in the timeseries as “swing” (forward velocity > 0 BL · s^−1^) or “stance” (forward velocity ≤ 0 BL · s^−1^) and find that our low-level locomotion clusters show signatures of distinct locomotory gaits (i.e., tetrapod and tripod gaits; Mendes et al. 2013; Pereira et al. 2019) with different numbers of legs being used for walking, on average, within each cluster. Together these results demonstrate that VAE-SNE is able to automatically decompose the dynamics of known complex behaviors (Video S1).

Due to the many philosophical complexities of objectively evaluating unsupervised cluster representations (reviewed by Jain et al. 1999; Kleinberg 2003; Todd et al. 2017), we forgo any further quantitative assessment of our clustering results and instead leave this for future work. For example, it is unclear how to best select the number of clusters for many different algorithms; how to properly compare algorithms that naturally produce different numbers of clusters and cluster shapes; and what metric(s) should be used to meaningfully evaluate a clustering description as generally “good” or “useful” other than manual, qualitative validation of the results, which we already provide here — though several quantitative descriptors with varying levels of desirability have been recently proposed for behavioral data (Todd et al., 2017). Comparing unsupervised cluster labels with a priori-defined labels — as is common practice (e.g., Jiang et al. 2016; Xie et al. 2016; Guo et al. 2017; Yang et al. 2019; Luxem et al. 2020) — is also problematic, as human-supervised descriptions may not accurately capture the underlying structure of the data distribution, and this is especially true for datasets where the goal is to potentially discover subtle differences that are undetectable by humans (e.g., Wiltschko et al. 2015). Despite the limitations imposed by these complexities, our results still illustrate multiple useful features of VAE-SNE as a general-purpose method.

Overall, we demonstrate how VAE-SNE can be used as a practical, scalable, and flexible tool for clustering real-world high-dimensional data. In this case, we transform posture data into interpretable behavioral labels that are comparable to those from previous methods (Berman et al., 2014, 2016; Todd et al., 2017; Klibaite et al., 2017; Cande et al., 2018; Klibaite and Shaevitz, 2019; Pereira et al., 2019). However, in contrast to many of these existing methods, VAE-SNE performs dimension reduction and clustering simultaneously, and unlike most previously-described algorithms for clustering data (e.g., Jiang et al. 2016; Xie et al. 2016; Guo et al. 2017; Yang et al. 2019), our method learns a small set of decipherable classes without the need to carefully tune the number of clusters fitted to the data, which can often be a non-trivial, unintuitive, and computationally-intensive process (Milligan and Cooper, 1985; Pham et al., 2005; Fang and Wang, 2012; Todd et al., 2017). Instead, any arbitrarily large number will give similar results due to the sparse watershed assignment procedure we use to combine overlapping clusters (Methods; Fig. 1d; Todd et al. 2017). In contrast to methods that impose strong assumptions about cluster shape, our clustering method has relaxed assumptions and allows for arbitrarily complex (e.g., non-convex) cluster distributions based on the local structure of the data. Additionally, in comparison to prior methods for unsupervised behavioral analysis, VAE-SNE has the advantage of being able to use more than two dimensions for clustering data, which has been shown to provide higher-quality behavioral labels with many potentially-desirable properties (Todd et al., 2017). Finally, our results further show that there is no need to carefully select a subset of data to use for training (e.g., the importance sampling technique described by Berman et al. 2014), which can also be a time-consuming process. Instead, VAE-SNE can be readily applied to large datasets that cannot fit into memory while still successfully detecting relatively short-lived and infrequent types of behavior, such as wing movements (Fig. 3b-c;Video S5).

## 3 Discussion

Here we introduce VAE-SNE, a deep generative model for simultaneously reducing dimensionality and clustering data. We compare VAE-SNE to existing methods for dimensionality reduction and demonstrate its utility and versatility using real-world examples. Our results establish that VAE-SNE is able to generate robust and interpretable compressed representations for data from different domains and is comparable in performance to other nonlinear methods for dimensionality reduction. In contrast to these existing methods, VAE-SNE has the advantage of being able to automatically cluster similar observations into a small set of classes, which can then be used to summarize large datasets with a coarse-grained description or select specific subpopulations of data for more detailed analysis. Our approach can also readily scale to very large datasets by leveraging techniques from deep learning — including, and especially, out-of-core data that cannot fit into memory. However, despite these strengths, VAE-SNE still has important limitations depending on the goals of the user, and there are many ways in which the model could be improved or extended in subsequent iterations. There are also other domains that VAE-SNE could be applied to in the future.

VAE-SNE preserves local relationships while also minimizing global structure distortion. Additionally, while VAE-SNE is not explicitly an autoregressive model, it still preserves a good deal of high-dimensional timeseries information. However, our results also show that VAE-SNE, and most of the other dimension reduction methods we tested, does not accurately preserve fine-scale structure (neighborhoods <1% of the total embedding size). For many applications, preserving these details may be unimportant, but this structure has been shown to be useful for detecting infrequent types of data, such as rare cell types (Linderman et al., 2019). Therefore, our results suggest that if researchers wish to preserve this type of information they should use FIt-SNE (Linderman et al., 2017, 2019) or Barnes-Hut-SNE (van der Maaten, 2014) over other algorithms for dimension reduction. We also find that, when initialized with PCA over the default initialization, UMAP (McInnes et al., 2018) preserves global structure slightly better without noticeably affecting local structure preservation, so PCA may be a more advantageous choice for initializing UMAP embeddings.

VAE-SNE optimizes faster than existing deep learning methods for dimensionality reduction, but FIt-SNE (Linderman et al., 2017, 2019), Barnes-Hut-SNE (van der Maaten, 2014), and UMAP (McInnes et al., 2018) are still faster. However, the training time for deep-neural-network methods like VAE-SNE and ivis (Szubert et al., 2019) can be variable due to the use of early stopping criteria that automatically end training when no improvement in the objective function is detected. These early stopping criteria could be easily adjusted to further shorten (or lengthen) training time. While we did not assess performance during the optimization process, much of the training time for VAE-SNE is spent on minor improvements to the objective function, which indicates adequate results can also be achieved with less training time. Additionally, FIt-SNE (Linderman et al., 2017, 2019), Barnes-Hut-SNE (van der Maaten, 2014), and UMAP (McInnes et al., 2018), are much slower for embedding new data because they calculate nearest neighbors for the new data and further optimize the embedding, which VAE-SNE does not require due to its learned encoder function. For smaller datasets that can fit in memory FIt-SNE (Linderman et al., 2017, 2019), Barnes-Hut-SNE (van der Maaten, 2014), and UMAP (McInnes et al., 2018) are still attractive options for dimensionality reduction, but for datasets that do no fit into memory, VAE-SNE provides some distinct advantages.

VAE-SNE has the ability to detect outliers and assess the embedding quality for out-of-sample data. This provides a straightforward mechanism for identifying new data to include in the training set, which can further improve performance. Most of the other algorithms we tested, or at least the specific software implementations we tested, provide no mechanism for quantitatively assessing embedding quality for each observation — with outliers being simply embedded under the assumption that the data are well supported by the training distribution. This can cause problems for any downstream analysis, especially when using statistical tests to answer scientific questions. Further improvements for outlier detection might include the use of Bayesian inference (Hafner et al., 2018) or other methods for estimating predictive uncertainty (reviewed by Kendall and Gal 2017).

We demonstrate that results produced by VAE-SNE can serve as a highly-interpretable coarse-grained description of tens-of-millions of observations — with several advantages over existing methods for clustering data. Applying VAE-SNE to future research in the behavioral sciences could help to reveal the genetic, environmental, and neural underpinnings of animal behavior (Berman, 2018; Brown and De Bivort, 2018; Datta et al., 2019) — especially when combined with recent advances in behavioral measurement (Mathis et al., 2018; Pereira et al., 2019; Graving et al., 2019; Günel et al., 2019) as well as genetic (Ran et al., 2013; Doudna and Charpentier, 2014), sensory (Stowers et al., 2017), and neural (Bath et al., 2014; Cande et al., 2018) manipulations. The clustering capabilities of VAE-SNE could also be applied to other types of data, such as single-cell RNA-seq data (Ding et al., 2018; La Manno et al., 2018) and natural history images (Cuthill et al., 2019; Zhang et al., 2019), but we leave this as future work for other researchers and domain experts to explore and validate. VAE-SNE might also be further improved by the use of more complex hierarchical clustering distributions (Tomczak and Welling, 2017; Roberts et al., 2018; Razavi et al., 2019), where additional scales with finer- or coarser-grained descriptions can be selected from the model for post-hoc analysis. Recent work has also shown that iteratively adjusting the parameters of the t-SNE similarity kernel can be used to generate a hierarchy of clusters in the latent embedding (Robinson and Pierce-Hoffman, 2020), which could be potentially applied to VAE-SNE as well.

To demonstrate the flexibility of VAE-SNE as a deep learning model, we introduce a variant for embedding data in polar coordinates on a unit sphere (Appendix C.1). We find that VAE-SNE successfully preserves structure in a spherical embedding as well (Fig. S10; Video S8;Video S9), which may be a more natural way to model some high-dimensional data sets (Davidson et al., 2018) since it avoids the “crowding” problem common to other embedding methods (van der Maaten and Hinton, 2008; Ding and Regev, 2019). While we focus on the Euclidean and cosine distances for calculating local neighborhoods, any differentiable distance function could potentially be substituted to create different embedding geometries, and, while we focus on kernels from the location-scale family of probability distributions (i.e. Gaussian, Student’s *t*), other log probability functions could potentially be used as well.

We also introduce a convolutional version of VAE-SNE for embedding images directly from raw pixel data (Appendix C.2). After applying this model to natural history images, we find that it groups perceptually-similar images based on complex sets of image features that correspond with taxonomic groupings (Figs. S11, S12). These results indicate that convolutional VAE-SNE may be useful for tasks such as relating distributions of complex animal coloration patterns to ecological, evolutionary, and behavioral function (Cuthill et al., 2017, 2019; Ezray et al., 2019; Wham et al., 2019). Future applications might include applying VAE-SNE to audio data (e.g., Oord et al. 2016; Sainburg et al. 2019).

There are multitude of ways in which VAE-SNE could be further improved or extended. Naturally, future work could apply more recent advances in variational and probabilistic inference like normalizing flows (Rezende and Mohamed, 2015; Kingma et al., 2016; Papamakarios et al., 2017), which allow data to be modeled with a more direct invertible mapping from the latent posterior to the data distribution, while also employing flexible, arbitrarily-complex distributions. The latent distribution used for VAE-SNE could also be modeled using many other types of representations such as quantized (Van Den Oord et al., 2017) or categorical (Jang et al., 2016; Maddison et al., 2016) distributions. Recent progress in generative adversarial networks (GANs; Goodfellow et al. 2014), may also provide further enhancements for modeling complex feature dependencies within the data distribution (Larsen et al., 2016; Srivastava et al., 2017; Dieng et al., 2019b). Timeseries data could be explicitly modeled using autoregressive deep neural networks (e.g., Oord et al. 2016) for the encoder and decoder similar to Wiltschko et al. (2015); Johnson et al. (2016b); Sussillo et al. (2016); Markowitz et al. (2018); Pandarinath et al. (2018); Luxem et al. (2020), and the latent distribution can be optimized to accurately predict future observations, which has been shown to be a useful framework for modeling behavior (Berman et al., 2016; Luxem et al., 2020). Additionally, computational efficiency might be further improved by applying recent advances in metric (Sohn, 2016) and contrastive learning (Chen et al., 2020), which may reduce or eliminate the need to perform expensive pairwise computations. Recent work on density-preserving versions of t-SNE and UMAP (Narayan et al., 2020) could also be incorporated to further improve the embedding quality.

Explicitly modeling hierarchical structure caused by variance across individual trials and subjects (Pandarinath et al., 2018) and batch effects due to variance in sampling procedures (Ding and Regev, 2019) is also important for improving VAE-SNE in the future. These effects could be accounted for with more complex, hierarchically-parameterized models (Sussillo et al., 2016; Pandarinath et al., 2018), hierarchical latent distributions (Tomczak and Welling, 2017; Roberts et al., 2018; Razavi et al., 2019), and new similarity kernels — such as the conditional t-SNE kernel recently proposed by Kang et al. (2019). The general use of conditional (e.g., Van den Oord et al. 2016) or supervised (e.g., Alemi et al. 2016) labels when optimizing the model could also help to integrate additional prior information about the data distribution into the latent distribution, the latter of which is already a feature of both UMAP (McInnes et al., 2018) and ivis (Szubert et al., 2019).

In summary, VAE-SNE is a general-purpose deep learning model for both dimension reduction and clustering that can be applied to many different types of data and readily scales to large datasets. Together our results illustrate that it is a robust, feature-rich method with multiple distinct advantages that make it an effective tool for analyzing real-world datasets across disciplines.

## 4 Methods

### 4.1 The VAE-SNE model

VAE-SNE is a variational autoencoder (VAE; Appendix A.1) with a learned Gaussian mixture prior (Kingma et al., 2014; Dilokthanakul et al., 2016; Tomczak and Welling, 2017) that is optimized using the ELBO objective function (derived in Appendix A.2) with an additional local neighborhood regularizer (Hinton and Roweis, 2003; van der Maaten and Hinton, 2008; van der Maaten, 2009; Ding et al., 2018). The likelihood and divergence terms from the ELBO objective can be broadly considered as an information theoretic trade-off between reconstruction accuracy (distortion) and compression (rate) respectively (Alemi et al., 2016; Chalk et al., 2016; Alemi et al., 2017), which makes VAEs an attractive solution for dimensionality reduction. However, there are implicit problems with the ELBO objective (reviewed by Alemi et al. 2017; Dieng et al. 2019a) that may prevent the model from learning a useful latent representation — e.g., a powerful, overparameterized decoder can simply ignore the compressed latent codes but still produce high-quality reconstructions. These issues render VAEs problematic as a general method for reducing dimensionality, as the primary purpose of dimensionality reduction is to create compressed representations that preserve important statistical features of the original data distribution.

#### 4.1.1 Regularizing the ELBO to improve structure preservation

We address the problems outlined above by optimizing VAE-SNE with a regularized version of the ELBO. This modification introduces a pairwise similarity regularizer derived from the (*t*-distributed) stochastic neighbor embedding (SNE/t-SNE) objective (Hinton and Roweis, 2003; van der Maaten and Hinton, 2008; van der Maaten, 2009). This idea of using the SNE objective for regularizing the latent space of VAEs was first proposed by Chien and Hsu (2017), which they called variational manifold probabilistic linear discriminant analysis (vm-PLDA), and later independently proposed by Ding et al. (2018) with their scvis model. However, the idea of applying the SNE objective to autoencoders, and deep neural networks in general, was introduced much earlier by van der Maaten(2009) with parametric t-SNE (pt-SNE), who proposed to use this objective in conjunction with an autoencoder to jointly learn a latent embedding. The pt-SNE model (van der Maaten, 2009) was also recently combined with advances from the Barnes-Hut-SNE algorithm (van der Maaten, 2014) under the name net-SNE (Cho et al., 2018). Additionally, Moody (2017) developed one of the first publicly-available pieces of software to combine the SNE objective with variational inference (variational t-SNE, or vt-SNE; and topic-SNE) but did not use a deep neural network to amortize inference across a set of shared parameters. Im et al. (2018) also proposed a variational bound on the t-SNE objective to improve optimization.

Here we apply the SNE objective to a VAE in a similar fashion to Ding et al. (2018). That is, we use the SNE objective as a method of better preserving structure in the latent embedding produced by our VAE, which improves the usefulness of the compressed representation (approximate posterior) produced by the ELBO. When combined into a single objective, we call this the stochastic neighbor evidence lower bound, or SNELBO. Generalizing from Ding et al. (2018), given a high-dimensional data matrix **X** = {**x**_1_, …, **x**_*N*_} and model parameters {***θ***, *ϕ*}, the SNELBO objective is written as:

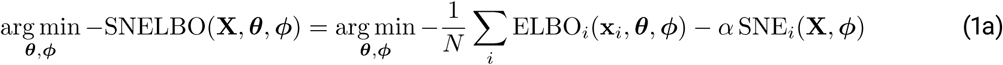

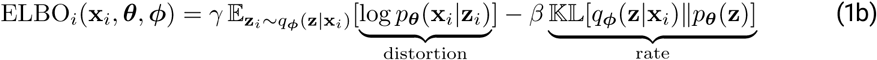

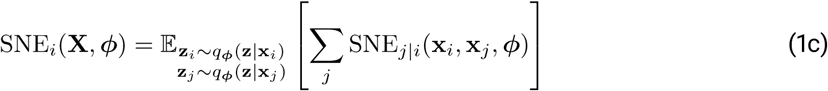

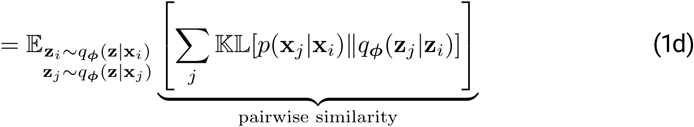

for *i, j* = 1, …, *N* and *i* ≠ *j*, where *N* is the number of observations in the *N* × *M* matrix **X** ∈ ℝ^*M*^. Thus vectors **x**_*i*_ and **x**_*j*_ are the *i*th and *j*th row in **X**, while **z**_*i*_ and **z**_*j*_ are Monte Carlo samples from the approximate low-dimensional posterior **z**_*i*_ ∼ *q*_*ϕ*_(**z**|**x**_*i*_) and **z**_*j*_ ∼ *q*_*ϕ*_(**z**|**x**_*j*_) respectively (Eq. 12c) — sampled using the reparameterization trick from Kingma and Welling (2013), or **z**_*i*_ = ***µ*** + ***σ*** ⨀ *ϵ*, where *ϵ* is an auxillary noise variable *ϵ* ∼ *N*(0, **I**) and ⨀ is the element-wise product (see Appendix A.3 for further discussion).

The objective function (Eq. 1a) consists of three terms, which can be interpreted as follows: (**1**) the expected log likelihood of the decoder distribution (Eq. 1b; distortion) minimizes distortion between the observed ground truth **x**_*i*_ and reconstruction, or maximizes accuracy, and preserves global structure in the embedding; (**2**) the divergence between the approximate posterior and the prior distribution (Eq. 1b; rate) constrains the global coordinate space of the embedding and restricts the rate of information (relative to the prior) that can be transmitted through the compressed space; and (**3**) the expected divergence between pairwise similarities (Eq. 1d) in high-dimensional space *p*(**x**_*j*_|**x**_*i*_) and those in low-dimensional space *q*_*ϕ*_(**z**_*j*_|**z**_*i*_) acts as a regularizer to preserve local neighbor relationships between data points. Further details of this stochastic neighbor regularizer are derived in Appendix B.

The Lagrange multipliers *γ, β*, and *α* are used to weight the distortion, rate, and pairwise similarity terms respectively, which we include as hyperparameters for the model. These multipliers can be adjusted to produce different forms of the objective for optimizing the model — e.g., increasing or decreasing the rate with the *β* multiplier (Higgins et al., 2017) — but in practice we set *γ* = *β* = 1, while *α* is set (following Ding et al. 2018) to the dimensionality of the data *α* = *M* to match the distortion term, which scales with the size of the input, or 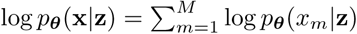.

#### 4.1.2 Learning a Gaussian mixture prior

For optimizing the VAE-SNE objective (Eq. 1a), we use a learned, or empirical, Gaussian mixture prior for *p*_*θ*_(**z**) which allows for an arbitrarily complex distribution (similar to Kingma et al. 2014; Dilokthanakul et al. 2016; Tomczak and Welling 2017). Using a more complex distribution allows for a tighter bound on objective, and, after optimization, approaches the true posterior distribution as the complexity of the distribution is increased (Kingma et al., 2014; Dilokthanakul et al., 2016; Tomczak and Welling, 2017; Cremer et al., 2017). The Gaussian mixture distribution is written as the weighted mixture of *K* Gaussian components:

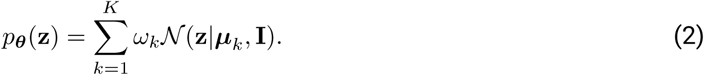

The mean ***µ***_*k*_ ∈ **M** and mixture weight *ω*_*k*_ ∈ ***ω*** of each component are learned as model parameters {**M, *ω***} ∈ ***θ*** subject to a softmax normalization constraint 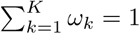. We also regularize the prior distribution by minimizing the divergence between the mixture distribution used to weight each component and a maximum-entropy mixture distribution, or:

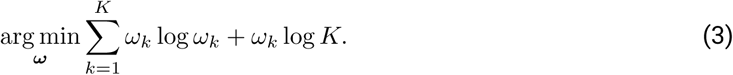

This prevents the prior from degenerating to a small number of modes (a problem described in more detail by Kingma et al. 2014; Dilokthanakul et al. 2016) by increasing the entropy of the mixture distribution. A higher entropy mixture distribution forces to model to utilize more of the components within the distribution, which increases the number of clusters and, consequently, the level of detail of the final clustering description (Still and Bialek, 2004). An analogous maximum entropy regularizer was also recently applied to solve the long-standing mode collapse problem common to generative adversarial networks (GANs; Dieng et al. 2019b).

The covariance for each component distribution could be learned as free parameters, but we find that using a simpler identity covariance matrix **I** allows for a sufficiently expressive prior distribution without adding additional complexity — and is less prone to cluster degeneracy during optimization. Using a highly-flexible (i.e., *K* ≫ 1) learned distribution as the prior for the latent space allows for better structure preservation, as non-convex structures are not distorted by the use of an overly simple prior. Also note that the special case of *K* = 1 mixture component is equivalent to the standard VAE prior (Kingma and Welling, 2013), or *p*_*θ*_(**z**) = *N*(**z**|0, **I**), which is the prior used by Ding et al. (2018).

##### Calculating the rate loss term

The parameters for the Gaussian mixture prior {**M, *ω***} ∈ ***θ*** are then learned from the data via the rate term in the VAE-SNE objective (Eq. 1b). For the special case of *K* = 1 we compute the Kullback-Leibler divergence analytically; however, because there is no analytical solution for a Gaussian mixture distribution with *K* > 1, we instead approximate this term numerically using Monte Carlo integration. In this case we use the expected log-density ratio for calculating the rate (Appendix A.2), which is written as:

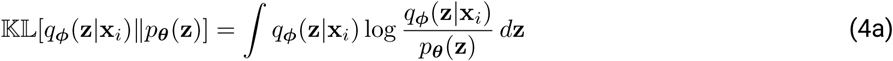

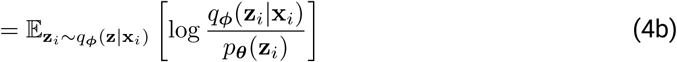

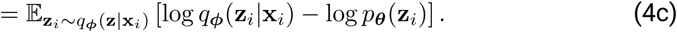

##### Clustering data with the Gaussian mixture prior

After optimizing the parameters for the prior, we can then use the learned Gaussian mixture to assign embedded data to discrete clusters. In other words, we wish to calculate the conditional distribution *p*_*θ*_(**y**|**z**), where **y** is a vector of class labels, or **y** = {*y*_1_, *y*_2_, …, *y*_*K*_}. However, the Gaussian mixture prior can contain highly-overlapping component distributions, which can cause undesirable side-effects. On one hand, this renders the parameterized mode for each overlapping component an unreliable descriptor of the surrounding local density, as each component is then simply a degenerate sub-mode within a non-Gaussian density cluster rather than a distinct subpopulation within the distribution delineated by the structure of the data. On the other hand, a Gaussian mixture distribution can have any arbitrary arrangement of weighted components, which makes the task of directly calculating the true local density mode for each embedded point both analytically and numerically intractable. Therefore, to circumvent these problems, we apply the sparse watershed assignment procedure described by Todd et al. (2017) to find the true local maximum for each component in the distribution — rather than for every embedded observation — through numerical optimization, which requires only a nominal amount of additional computation. We can then merge overlapping components and assign embedded data to a mode that more accurately reflects the underlying (potentially non-Gaussian) region of local density.

Because this sparse watershed procedure produces clusters with an arbitrary number of weighted components, calculating the full posterior probability *p*_*θ*_(**y**|**z**) for each data point is computationally complex. So for the sake of simplicity, we perform hard label assignment. In other words, we calculate the mode of the cluster distribution for each value of **z**, or:

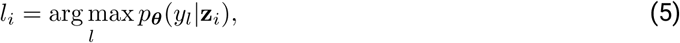

for *l* = 1, …, *K*, where *l*_*i*_ is the assigned label for the latent vector **z**_*i*_. This hard label assignment procedure is performed in 3 steps: (**1**) latent vectors are initially assigned to the nearest (highest local density) component in the Gaussian mixture prior; (**2**) the Gaussian mixture distribution is further optimized to combine overlapping mixture components using sparse watershed assignment (Todd et al., 2017); and (**3**) the initial cluster assignments are then recursively updated using the learned hierarchy of overlapping components to ensure each latent vector is assigned to the mode that best represents the underlying density of the local neighborhood for that observation. To accomplish these steps, the expected value of the approximate posterior for each data point is initially assigned to a single mode in the Gaussian mixture distribution by calculating the weighted mixture component with the maximum likelihood (minimum distortion), which is written as:

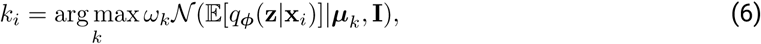

where *k*_*i*_ is the initial cluster assignment for the *i*th data point **x**_*i*_. We then combine degenerate (highly-overlapping) modes from the distribution by applying the sparse watershed procedure described by Todd et al. (2017). Using this procedure, the initial cluster assignments are further combined by optimizing the mean of each component to ascend to its local maximum within the Gaussian mixture prior, which we write as a minimization of the negative log-likelihood, or:

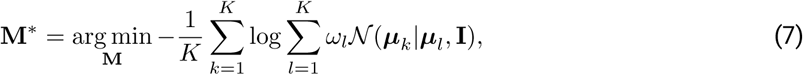

where 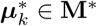 is the optimized mean of each component. We optimize this objective numerically with the Adam optimizer (Kingma and Ba, 2014) with a learning rate of 1 × 10^−3^ until the objective (Eq. 7) stops improving for 100 training steps. We then merge cluster assignments based on whether the mode for the initial cluster assignment *k*_*i*_ has moved within the basin of attraction for another mixture component in the distribution (after optimizing Eq. 7), or:

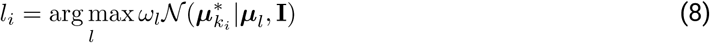

where *l*_*i*_ is the sparse watershed label assignment for the *i*th data point **x**_*i*_, which was assigned to the *k*_*i*_th mode of the distribution 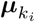 in the initial cluster assignment step (Eq. 6). We then repeat this assignment procedure *K* times to ensure all label assignments to degenerate modes are reassigned to the mode with the highest local density:

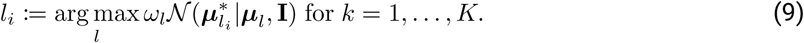

Note that, for data assigned to non-degenerate modes in the initial step, typically the cluster assignment remains unchanged, where *l*_*i*_ = *k*_*i*_.

### 4.2 Comparing dimensionality reduction algorithms

We compared VAE-SNE to other dimensionality reduction algorithms including PCA (scikit-learn v0.23.0; Pedregosa et al. 2011), t-SNE (van der Maaten and Hinton, 2008), UMAP (v0.4.0; McInnes et al. 2018), scvis (Ding et al., 2018), and ivis (v1.7.2; Szubert et al. 2019). Our main comparisons involve compressing data to two dimensions for visualization purposes, but VAE-SNE (and other algorithms) can be used for dimensionality reduction more generally.

#### 4.2.1 openTSNE and t-SNE variants

For t-SNE we used the openTSNE (v0.4.0) implementation from Poličar et al. (2019), which includes improvements from van der Maaten (2014); Linderman et al. (2017, 2019) to maximize speed and scalability, as well as methods for embedding out-of-sample data described by Poličar et al. (2019) (see also Berman et al. 2014; Kobak and Berens 2019). We tested two versions of openTSNE using both the Barnes-Hut approximation (Barnes-Hut-SNE) from van der Maaten (2014) and the Fourier interpolation approximation (FIt-SNE) from Linderman et al. (2017, 2019). However, FIt-SNE, the fastest version of openTSNE, is practically limited to very low dimensional embeddings (i.e., 1-D or 2-D) due to the Fourier interpolation algorithm used for approximating the gradient during optimization, and therefore cannot be used for more general-purpose dimensionality reduction (Linderman et al., 2017, 2019).

#### 4.2.2 scvis as a special case of VAE-SNE

We found the original implementation of scvis (Ding et al., 2018) difficult to use for our comparisons without extensive modification, as it relies on outdated software dependencies and is limited to specific data file formats for using the code. However, scvis (Ding et al., 2018) can be considered a special case of VAE-SNE with specific hyperparameter settings, so instead we used VAE-SNE with hyperparameters matched to those described by Ding et al. (2018) for making comparisons. In particular, we used the network architecture for the encoder and decoder networks described by Ding et al. (2018), along with ELU activations (Clevert et al., 2015). We also use the asymmetric similarity kernel for the high-dimensional similarities (Eq. 17a), and we set *K* = 1 for the number of components in the prior distribution (Eq. 2). For benchmarking the processing speed of scvis (Ding et al., 2018), we disabled our added parallel computations (Section 4.5) to match the speed of the original implementation from Ding et al. (2018), and we calculated training time based on the original recommendation from Ding et al. (2018) for training with batch size of 512 for 100 epochs.

#### 4.2.3 Setting hyperparameters for comparisons

For each algorithm we used Euclidean distances for calculating pairwise similarities (the default for all of the algorithms tested) along with the default settings for all other hyperparameters with some exceptions. For t-SNE, we set n jobs=-1 to enable parallel processing. For UMAP, we also compare PCA initialization for the low-dimensional embedding (vs. the default Laplacian Eigenmap initialization), which is not a default option but improves global structure preservation. For ivis (Szubert et al., 2019), we used the default model and followed recommendations from Szubert et al. (2019) to adjust the early stopping criteria for different dataset sizes.

The hyperparameters for different methods could, of course, be adjusted ad infinitum to produce different types of embeddings and could bias performance for different datasets in many ways; however, the comparisons we make in this paper are not meant to be exhaustive, only informative in terms of validating VAE-SNE as a comparable method. In the end, researchers will have to decide for themselves which algorithm is most useful for their specific application. It is also worth considering that, for some of the algorithms tested, adjusting the hyperparameters can dramatically alter computational and memory requirements — for example, increasing the perplexity hyperparamater for FIt-SNE (Linderman et al., 2017) and Barnes-Hut-SNE (van der Maaten, 2014) or the `n_neighbors` hyperparameter for UMAP, increases number of nearest neighbors that are computed and, consequently, the size of the nearest neighbors graph used to optimize the embedding. Our decision to use default settings is also especially reasonable for the t-SNE variants we tested given that the openTSNE package (Poličar et al., 2019) uses hyperparameter suggestions from Kobak and Berens (2019), which have been empirically shown to work well across many datasets.

#### 4.2.4 VAE-SNE hyperparameters

We tested multiple variants of VAE-SNE in our comparisons, but across these variants we use similar hyperparameters for training. For the encoder and decoder networks we use 4 densely-connected layers each with 256 units (with biases). For each layer we apply the nonlinear SELU activation function and use the appropriate random initialization for the weights described by Klambauer et al. (2017). We train each VAE-SNE model for a maximum of 100 epochs with an initial batch size of 512 using the Adam optimizer (Kingma and Ba, 2014) with a learning rate of 0.001. For the perplexity hyperparameter, we calculate this as a function of the batch size used during training, which we call the *perplexity ratio*, such that P = *bϱ* where P is the perplexity, *b* is the batch size, and *ϱ* is the perplexity ratio. To improve global structure preservation, we begin training with *ϱ* = 0.1 and then anneal to *ϱ* = 0.01 by exponentially decaying *ϱ* after each training batch (similar to the perplexity annealing technique described by Kobak and Berens 2019). After the perplexity ratio is fully annealed to the target value, we then perform early stopping if pairwise similarity loss stops improving by at least 0.001 per epoch with a patience of 5 epochs (lack of progress is ignored for 5 epochs before stopping training). While it is common practice to decrease the learning rate after training stagnates to further improve performance, we instead increase the batch size, which has been shown to provide similar improvements (Smith et al., 2017). Therefore after training stagnates and early stopping is initiated for the initial batch size of 512, we increase the batch size to 1024 and continue training until early stopping is initiated again using the same criteria. For the Gaussian mixture prior we set the number of components to *K* = 100, but we found that any arbitrarily large number of components produced similar (nearly identical) results.

We tested 4 variants of VAE-SNE with different similarity kernels. We tested VAE-SNE using a t-SNE similarity kernel with (**1**) constant kernel parameters (*ν* = *τ* = 1) as well as (**2**) learned kernel parameters (van der Maaten, 2009). We also tested VAE-SNE variants using a SNE kernel with (**3**) constant (*η* = 1) and (**4**) learned parameters as well. Otherwise the hyperparameters for each variant were kept constant, as described above.

#### 4.2.5 Local structure preservation

After embedding the data with each algorithm we assessed local structure preservation with two measures of preservation that define local neighborhoods in different ways. For both of these metrics we targeted neighborhoods that correspond to ∼ 1% of the total embedding size.

##### metric-based neighborhoods

First, we used a metric-based measure of local neighborhood preservation, where neighborhoods are defined based on distance (a fixed radius) to a cluster center. Following Becht et al. (2019) we applied the k-means clustering algorithm (with *k* = 100 clusters; using scikit-learn v0.23; Pedregosa et al. 2011) to the high-dimensional data and the low-dimensional embedding for each method, which effectively divides the data into small Voronoi regions. We then calculated the normalized mutual information (reviewed by Vinh et al. 2010; see also McDaid et al. 2011) between the high-dimensional and low-dimensional cluster assignments (using scikit-learn v0.23; Pedregosa et al. 2011). This provides a symmetric and permutation invariant measure of how well local neighborhood memberships from the high-dimensional space are preserved by each embedding method — with similarity ranging from 0 (no overlap, or random) to 1 (perfect overlap). We performed 5 replicates of this for each trial.

##### topological neighborhoods

Second, we assessed local neighborhood preservation topologically by calculating the exact nearest neighbors for 1000 randomly selected data points and then defining the local neighborhood for each point as *k* nearest neighbors, where *k* is selected such that 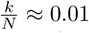, and *N* is the total embedding size. We then computed the proportion of the neighbors that are assigned to the correct local neighborhood in low-dimensional embedding, which ranges from 0 (no neighbors preserved) to 1 (all neighbors preserved). We performed 5 replicates of this for each trial.

#### 4.2.6 Global structure preservation

To assess global structure preservation we follow Becht et al. (2019) by calculating the Pearson correlation between pairwise squared Euclidean distances for 10,000 points in the high-dimensional space and the low-dimensional embedding for each method (for a total of 49.995 million distances). As distances have a lower bound of zero and tend to follow a log-normal (or Gamma) distribution, we first log transformed the distances in order to homogenize the variance and better match the assumptions of Pearson’s correlation score. The Pearson correlation then provides a measure of the global structure preservation ranging from -1 (anti-correlated) to 1 (correlated). We performed 5 replicates of this for each trial.

#### 4.2.7 Fine-scale structure preservation

Because our metrics for local structure preservation only account for a single scale but not the fine-scale structure within local neighborhoods, we also assessed topological structure preservation for smaller neighborhood sizes. As before, we calculated the exact nearest neighbors for 1000 randomly selected data points. We then computed the proportion of points assigned to the correct neighborhood across 14 dyadically (log_2_) spaced neighborhood sizes ranging from *k* = 2^1^ to *k* = 2^14^. Neighborhood sizes were then normalized as a proportion of the total embedding size, or 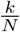. We performed 5 replicates of this for each trial and neighborhood size.

#### 4.2.8 Temporal structure preservation

Because the largest dataset we use is also timeseries data, we assess temporal structure preservation for the test set by calculating Euclidean distances between sequential time points in high-dimensions and low-dimensions for each method. We then calculate the Pearson correlation coefficient of the log transformed distances (same as for assessing global structure preservation) for 50 randomly selected 10 minute subsets (60,000 observations) within the full timeseries. This then provides a measure of how well temporal derivatives are preserved in the low-dimensional embedding ranging from -1 (anti-correlated) to 1 (correlated).

#### 4.2.9 Hierarchical bootstrap for statistical comparisons

To compare each information preservation metric statistically we performed hierarchical bootstrapping (see Saravanan et al. 2019 for a recent review). Every trial for each dimension reduction method has multiple observations per metric, which creates hierarchical dependencies in the data. To account for this, we use seaborn v0.10.1 (Waskom et al., 2020) to calculate and plot hierarchical bootstrap estimates of the mean for each information preservation metric — resampling (with replacement) both within trials and across trials (n=1000 bootstrap samples). We then plot the 95% intervals of the bootstrap distribution to compare the performance of each dimension reduction method statistically. Rather than attempting to make decisions regarding the statistical “significance” of these bootstrap distributions based on an arbitrary threshold, we instead simply treat them as a measure of the uncertainty (variance) in effect size for each information preservation metric. The computational experiments from which the information preservation metrics are derived could be run ad infinitum to achieve statistical significance, which is effectively a measure of statistical resolution based on the number of observations, but this is not necessarily informative in practice.

### 4.3 Datasets

#### 4.3.1 Animal body posture dynamics

The largest dataset we used for comparisons is a behavioral dataset from Berman et al. (2014, 2016); Pereira et al. (2019) consisting of ∼1-h video recordings (at 100Hz) for 59 freely-behaving individual fruit flies (*Drosophila melanogaster*) for a total of ∼21.1 million observations (downloaded from: http://arks.princeton.edu/ark:/88435/dsp01pz50gz79z). We tracked the full body posture of each individual with DeepPoseKit v0.3.6 (Graving et al., 2019) using the procedures described by Graving et al. (2019) to train a deep convolutional pose estimation model using the keypoint annotations from Pereira et al. (2019) as training data. For each video this produced a multivariate time series of the Euclidean coordinates describing 32 body part positions in the video — including the head, neck, eyes, thorax, abdomen, wings, and 24 leg joints. We then rotationally and translationally aligned the posture data at each timepoint to the major body axis (neck-thorax vector) and calculated the sine and cosine of the keypoint angles for the 30 body parts not used for alignment. This resulted in a 30 × 2 = 60 dimensional posture timeseries. To transform the spatial posture data into a dynamical spatio-temporal representation, we then applied a normalized Morlet wavelet transform from Berman et al. (2014) using the behavelet Python package v0.0.1 (Graving, 2019) to generate a multi-scale time-frequency spectrogram of the body posture dynamics for each time point. Following Berman et al. (2014); Pereira et al. (2019), we used 25 dyadically (log_2_) spaced frequencies ranging from 1Hz to 50Hz (the Nyquist frequency of the signal), which expanded the dimensionality of the timeseries from 30 × 2 = 60 to 30 × 2 × 25 = 1500.

##### Dimension reduction comparisons

To generate a training set for benchmarking the different algorithms, we uniformly randomly sampled a subset of data from the body posture dynamics timeseries for 58 of 59 individuals while excluding one randomly selected individual to use as a test set. We tested 4 training set sizes: 58 × 500 = 29, 000; 58 × 1000 = 58, 000; 58 × 2000 = 116, 000; 58 × 4000 = 232, 000, above which we encountered out-of-memory errors when running UMAP (McInnes et al., 2018) on larger subsets of data. Each test set contains ∼ 360, 000 sequential observations. We then applied each dimension reduction method to the training set and subsequently embedded the test set. For training VAE-SNE we used the cross-entropy loss as a log likelihood function, as it matches well with the normalized time-frequency data, but we also found that other likelihood functions work similarly well.

##### Behavioral clustering

To simplify the dataset for performing our clustering analysis, we used the sine and cosine of the keypoint angles for the 6 legs (the distal tips of each leg), 2 wings, head, and abdomen for a total of 10 body parts and a 10 × 2 = 20 dimensional posture timeseries. As before we applied the time-frequency transform which expands the dimensionality of the timeseries from 10 × 2 = 20 to 10 × 2 × 25 = 500. We then applied VAE-SNE with a t-SNE kernel (Appendix B; *ν* = *τ* = 1) to compress the spectrogram data to 30 dimensions. We used the cross-entropy between normalized time-frequency vectors, or ℍ[**x**_*i*_, **x**_*j*_] = ∑ **x**_*i*_ log **x**_*j*_, as our metric for calculating high-dimensional similarities (Appendix B), as this provides a more natural measure of divergence between the normalized spectrograms than Euclidean distance. The cross-entropy is closely related (up to a constant) to the Kullback-Leibler divergence — the metric originally used by Berman et al. (2014) — but is slightly faster to calculate, which reduces training time. When visualizing the spectrograms we integrate (sum) across the wavelet coefficients for the sine and cosine for each body part in the spectrogram.

#### 4.2.3 Single-cell RNA-seq

To test the application of VAE-SNE to single-cell RNA-seq data, we used data from La Manno et al. (2018) which consists of 18,213 observations describing the development and cell fate of hippocampal neurons. We preprocessed these data using the velocyto.py (v0.17.17) package from La Manno et al. (2018). We compressed the raw expression values to 500 dimensions using PCA before applying subsequent dimension reduction algorithms. We applied each dimension reduction algorithm to the full dataset and then re-embedded the training set in place of a test set in order to evaluate the speed for embedding new data. We report information preservation metrics only for the training set, as no test set was used due to the relatively small size of the dataset. For training VAE-SNE on this dataset we use a Student-t likelihood function, but found other likelihood functions work similarly well.

#### 4.3.3 Natural history images

We also applied a convolutional variant of VAE-SNE to natural history images, and to test this we used two datasets: a set of 59,244 shell images from Zhang et al. (2019) and a set of 2,468 butterfly images from Cuthill et al. (2019). All images were preprocessed by applying local adaptive thresholding to detect and remove the background. Images were then zero-padded to create a 1:1 aspect ratio and resized to a resolution of 192 × 192. We trained convolutional VAE-SNE using the same hyperparameters as the dimension reduction experiments, but using batches of only 256 images.

### 4.4 Computing hardware

All performance comparisons were conducted on a high-end consumer-grade workstation equipped with an Intel Core-i9-7900X CPU (10 cores, 20 threads @ 3.30GHz), 32GB of DDR4 RAM, a 4TB NVMe solid state drive, and a NVIDIA GeForce GTX 1080 Ti GPU (11 GB GDDR5X VRAM).

### 4.5 Parallelizing pairwise computations to improve performance

To improve performance of pairwise computations over Ding et al. (2018), we reimplemented the underlying algorithms for training VAE-SNE. The largest performance bottleneck for VAE-SNE is the recursive binary search algorithm for computing high-dimensional pairwise similarities (Appendix B). However, the computations for this algorithm are embarrassingly parallel, so we reimplemented it to run recursion loops in parallel across multiple CPU threads. This was accomplished by JIT-compiling the code using the numba library (Lam et al., 2015), which resulted in massive speed improvements. We also reimplemented all pairwise distance calculations on the GPU using PyTorch (Paszke et al., 2019), which further improved performance.

### 4.6 Code availability

The code for VAE-SNE is freely available at https://github.com/jgraving/vaesne under a permissive open-source license. The library is written primarily using PyTorch v1.5.0 (Paszke et al., 2019) and includes a scikit-learn-style API (Buitinck et al., 2013) for fitting the model (`model.fit()`) and predicting on new data (`model.predict()`).

## Acknowledgments

We thank Gordon Berman and members of the Berman lab, Tim Landgraf, Ben Wild, David Dormagen, Conor Heins, Blair Costelloe, Ian Etheredge, and Ari Strandburg-Peshkin for their helpful comments on the manuscript. We are also grateful to Mike Costelloe for his creative advice on the figures, Einat Couzin-Fuchs and Dan Bath for their expert input, and Michael Smith for the use of his GPU. J.M.G. and I.D.C. acknowledge support from the Deutsche Forschungsgemeinschaft (DFG, German Research Foundation) under Germany’s Excellence Strategy - EXC 2117 - 422037984. I.D.C. acknowledges support from NSF Grant IOS-1355061, Office of Naval Research Grant (ONR, N00014-19-1-2556), the Struktur-und Innovationsfonds für die Forschung of the State of Baden-Württemberg, and the Max Planck Society.

## Author Contributions

- **J.M.G**. — Conceptualization, Data curation, Software, Formal analysis, Validation, Investigation, Visualization, Methodology, Writing–original draft, Project administration, Writing–review and editing
- **I.D.C**. — Conceptualization, Resources, Writing–reviewing and editing, Supervision, Project administration, Funding acquisition

## Supplemental Figures

**Table S1.** Ranked information preservation metric performance for nonlinear dimension reduction algorithms. Rankings for each nonlinear dimension reduction algorithm in terms of general performance for local, global, fine-scale, and temporal structure preservation (lower is better).

| name | citation | local | global | fine-scale | temporal |
| --- | --- | --- | --- | --- | --- |
| VAE-SNE (t-SNE) | this paper | 1 | 1 | 2 | 2 |
| VAE-SNE (SNE) | this paper | 2 | 1 | 2 | 1 |
| Flt-SNE | Linderman et al. (2017) | 1 | 2 | 1 | 2 |
| Barnes-Hut-SNE | van der Maaten (2014) | 1 | 2 | 1 | 2 |
| UMAP (LE init) | McInnes et al. (2018) | 1 | 3 | 2 | 3 |
| UMAP (PCA init) | McInnes et al. (2018) | 1 | 2 | 2 | 3 |
| scvis | Ding et al. (2018) | 2 | 1 | 2 | 2 |
| ivis | Szubert et al. (2019) | 2 | 1 | 3 | 1 |

**Table S2.** Ranked processing speed performance for nonlinear dimension reduction algorithms. Rankings for each nonlinear dimension reduction algorithm in terms of general performance for training time and test time (lower is better), as well as whether or not test time increases as a function of training set size.

| name | citation | train time | test time | test time ∝ train size |
| --- | --- | --- | --- | --- |
| VAE-SNE | this paper | 4 | 1 | no |
| Flt-SNE | Linderman et al. (2017) | 2 | 4 | no |
| Barnes-Hut-SNE | van der Maaten (2014) | 3 | 5 | yes |
| UMAP | McInnes et al. (2018) | 1 | 3 | yes |
| scvis | Ding et al. (2018) | 6 | 1 | no |
| ivis | Szubert et al. (2019) | 5 | 2 | no |

**Table S3.** Additional features for nonlinear dimension reduction algorithms. A summary of potentially useful additional features for each nonlinear dimension reduction algorithm including batch training for applying dimension reduction to large out-of-core datasets, non-Euclidean embeddings for different types of compressed representations, whether the algorithm is tractable in higher dimensions (>2), and whether the algorithm learns a distribution of clusters within the data.

| name | citation | batch training | non-Euclidean | $>2$ dims. | clustering |
| --- | --- | --- | --- | --- | --- |
| VAE-SNE | this paper | yes | yes | yes | yes |
| Flt-SNE | Linderman et al. (2017) | no | no | no | no |
| Barnes-Hut-SNE | van der Maaten (2014) | no | no | yes | no |
| UMAP | McInnes et al. (2018) | no | yes | yes | no |
| scvis | Ding et al. (2018) | yes | no | yes | no |
| ivis | Szubert et al. (2019) | yes | no | yes | no |

**Figure S1.**
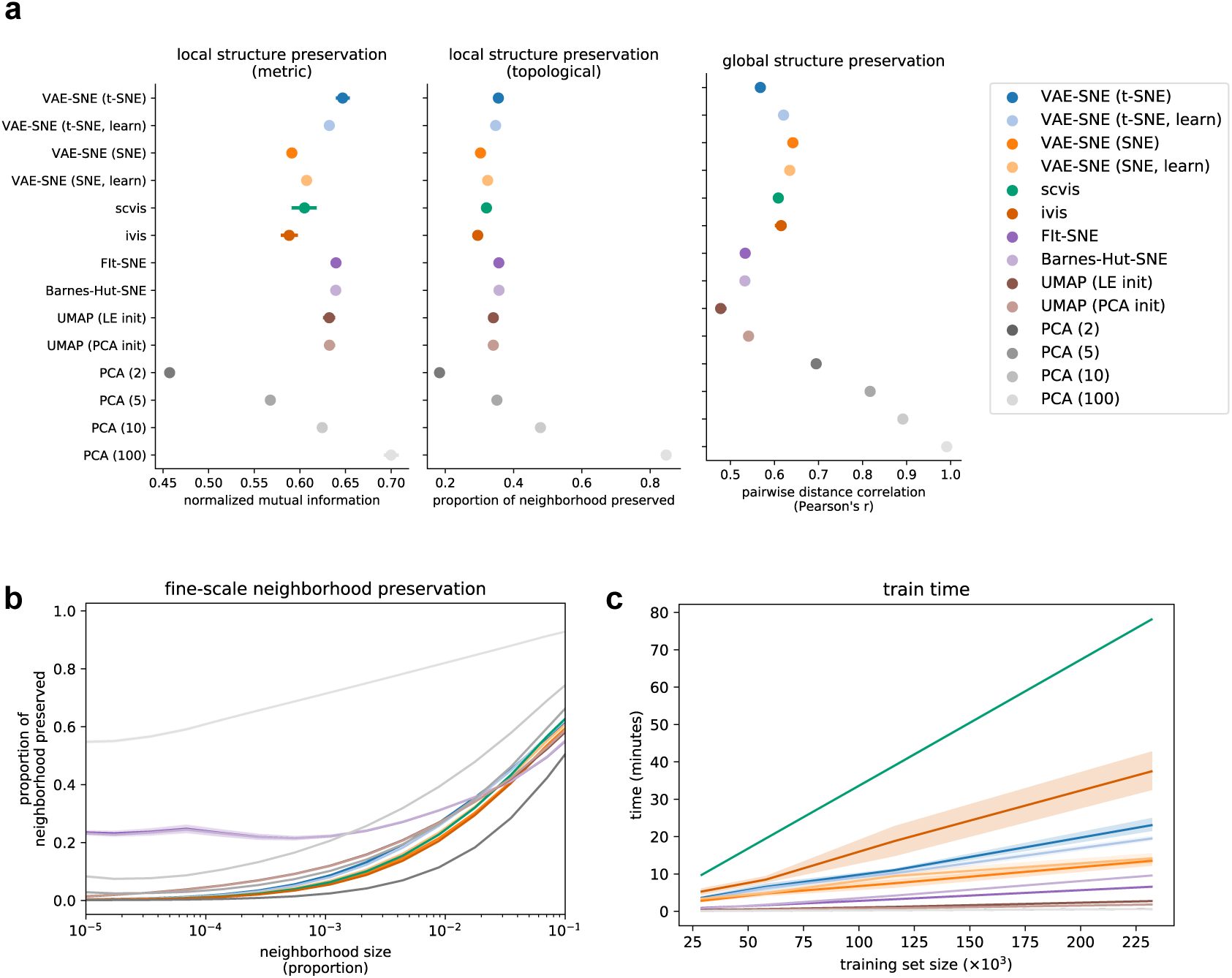
Dimension reduction performance for the posture dynamics training set. Plots show performance comparisons for the posture dynamics dataset (Berman et al., 2014, 2016; Pereira et al., 2019) using the training set. **a**, Mean and 95% interval of the bootstrap distribution for local and global structure preservation. Results are pooled across all training set sizes (for each metric n = 4 training set sizes × 5 trials × 5 replicates = 100 per algorithm). **b**, Mean and 95% interval of the bootstrap distribution for fine-scale structure preservation across multiple neighbor sizes (as a proportion of the total embedding size). Results are from the largest training set size only (n = 14 neighborhood sizes × 5 trials × 5 replicates = 350 per algorithm). **c**, Training time for fitting each algorithm across different training set sizes (n = 4 training set sizes × 5 trials = 20 per algorithm).

**Figure S2.**
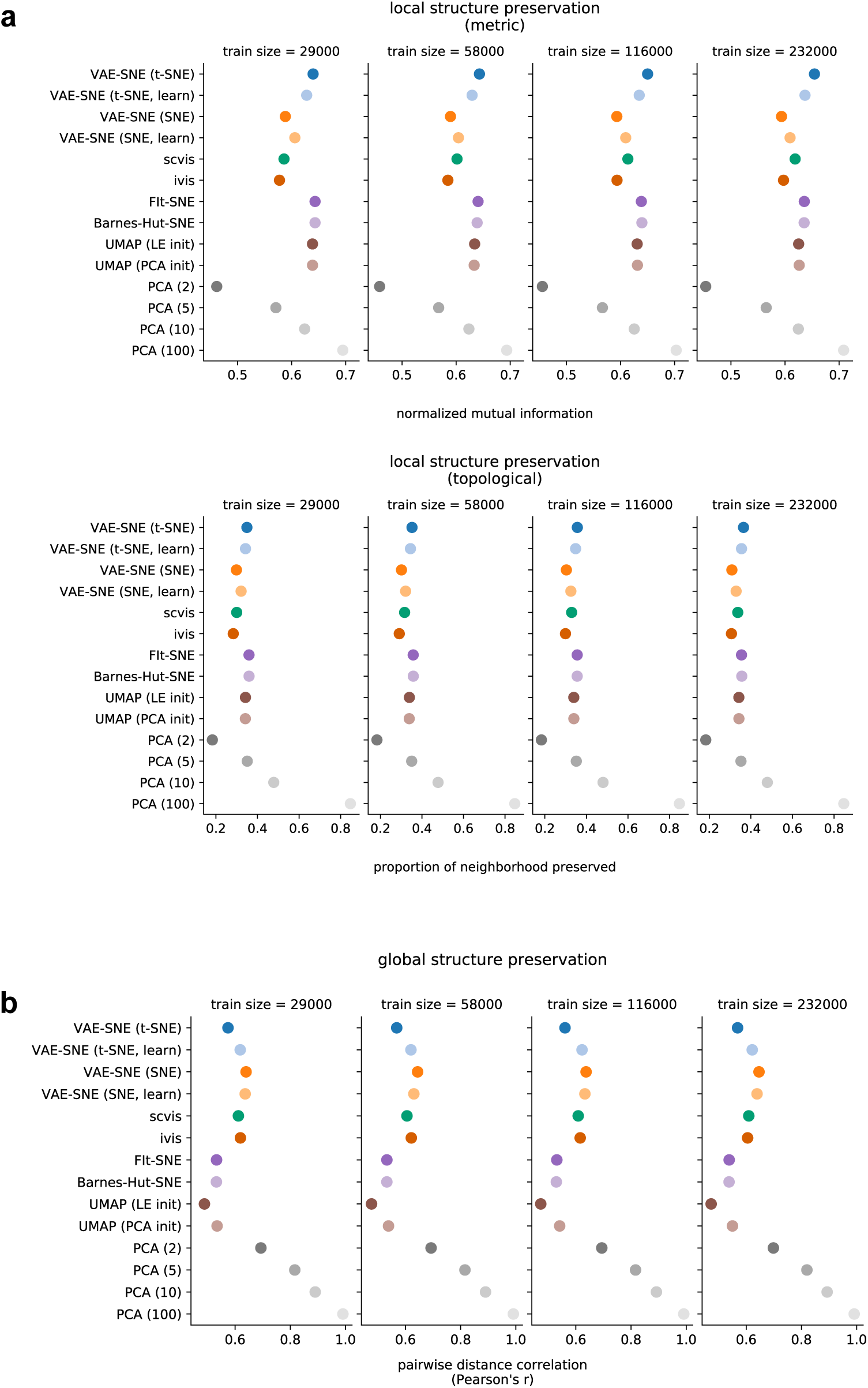
Dimension reduction performance for the posture dynamics training set across training set sizes. Plots show performance comparisons for the posture dynamics dataset (Berman et al., 2014, 2016; Pereira et al., 2019) using training sets of different sizes. **a-b**, Mean and 95% interval of the bootstrap distribution for local (**a**) and global (**b**) structure preservation. (for each metric n = 5 trials × 5 replicates = 25 per training set size per algorithm)

**Figure S3.**
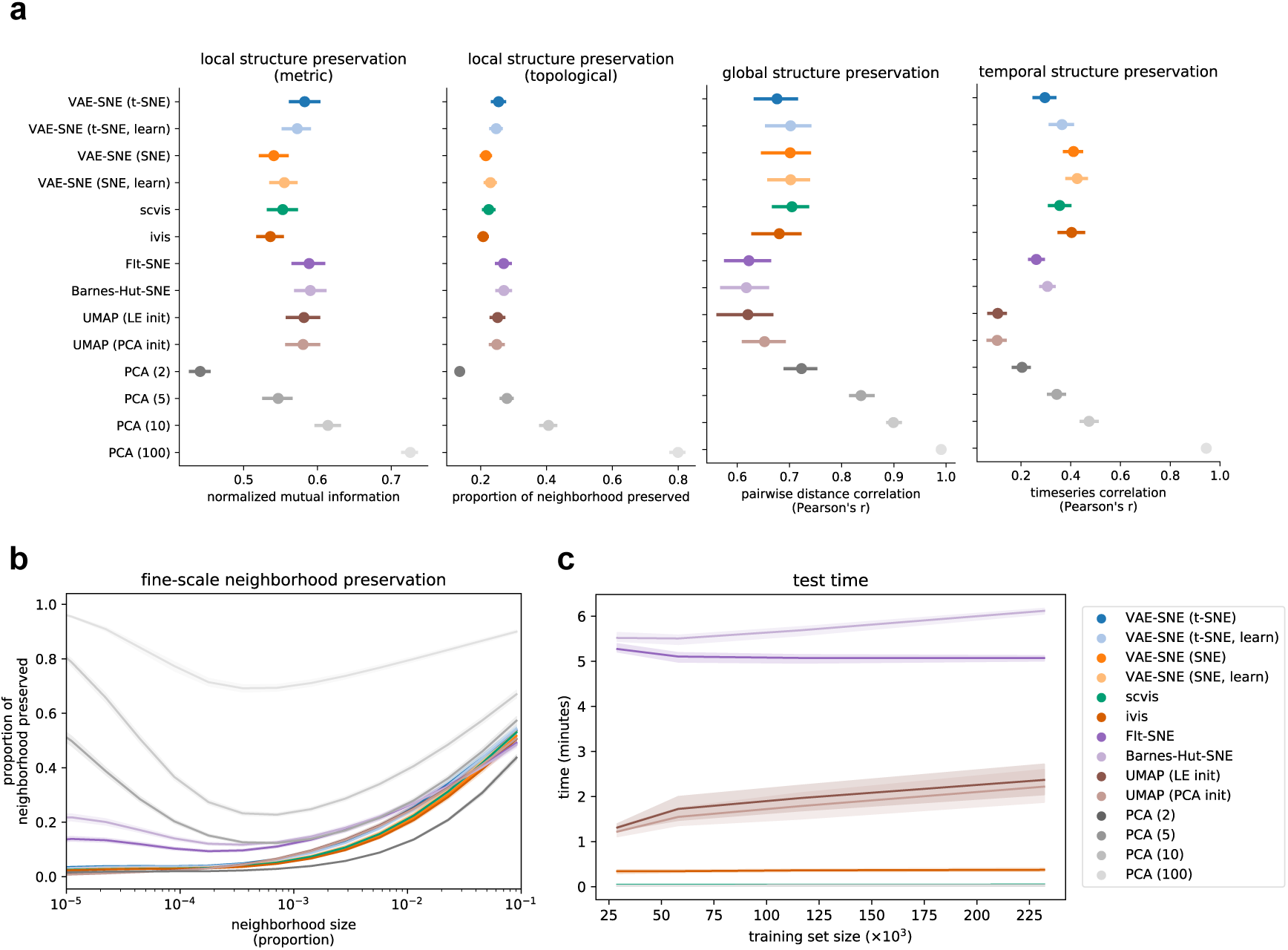
Dimension reduction performance for the posture dynamics test set. Plots show performance comparisons for the posture dynamics dataset (Berman et al., 2014, 2016; Pereira et al., 2019) using the test set. **a**, Mean and 95% interval of the bootstrap distribution for local, global, and temporal structure preservation. Results are pooled across all training set sizes (for local and global structure n = 4 training set sizes × 5 trials × 5 replicates = 100 per algorithm; for temporal structure n = 4 training set sizes × 5 trials × 50 subsamples = 1000 per algorithm). **b**, Mean and 95% interval of the bootstrap distribution for fine-scale structure preservation across multiple neighbor sizes (as a proportion of the total embedding size). Results are from the largest training set size only (n = 14 neighborhood sizes × 5 trials × 5 replicates = 350 per algorithm). **c**, Elapsed time for embedding the test set with each algorithm across different training set sizes (n = 4 training set sizes × 5 trials = 20 per algorithm).

**Figure S4.**
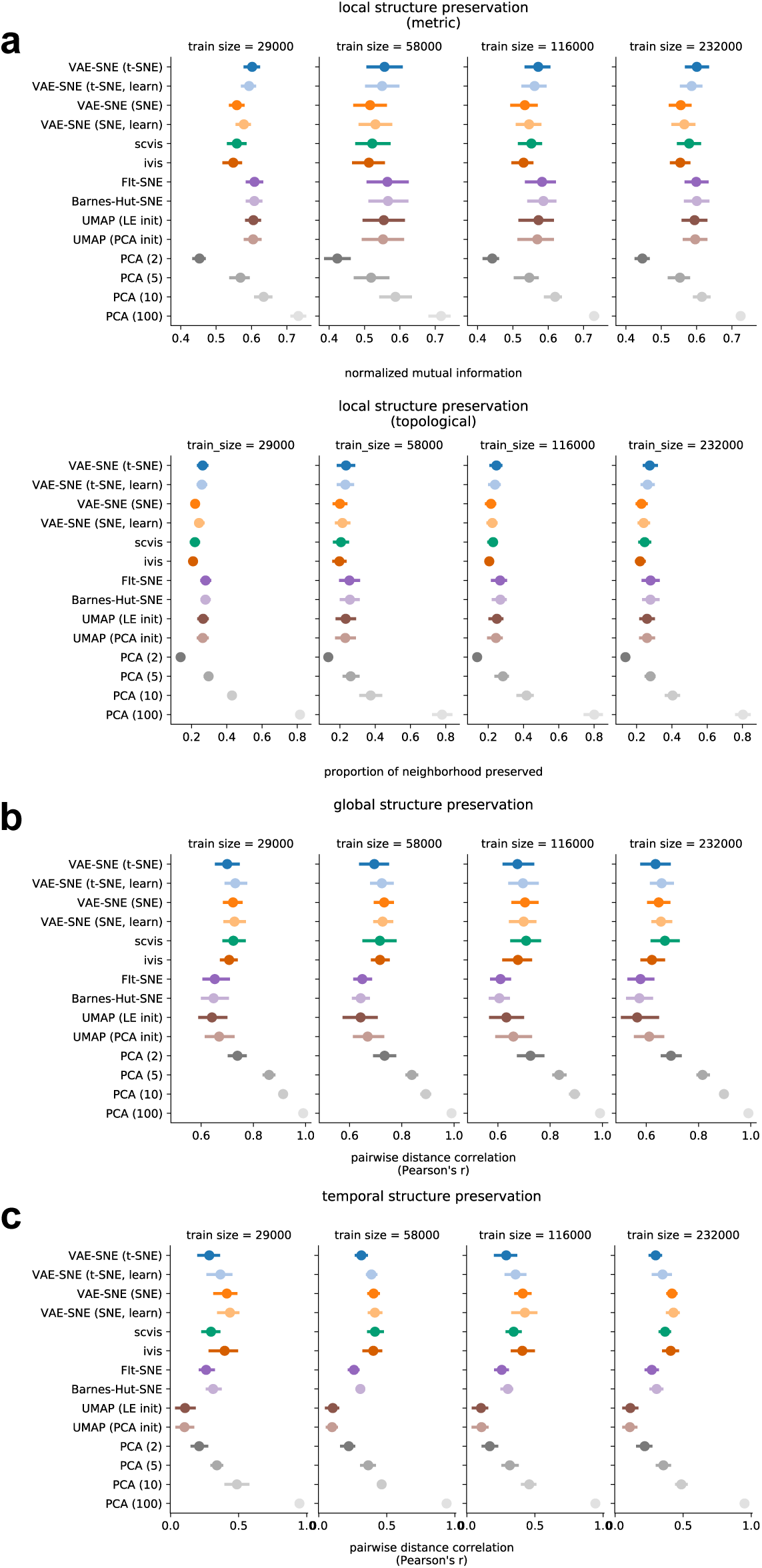
Dimension reduction performance for the posture dynamics test set across training set sizes. Plots show performance comparisons for the posture dynamics dataset (Berman et al., 2014, 2016; Pereira et al., 2019) using training sets of different sizes. **a-c**, Mean and 95% interval of the bootstrap distribution for local (**a**), global (**b**), and temporal (**c**) structure preservation (for local and global structure n = 5 trials × 5 replicates = 25 per algorithm for each training set size; for temporal structure n = 5 trials × 50 subsamples = 250 per algorithm for each training set size).

**Figure S5.**
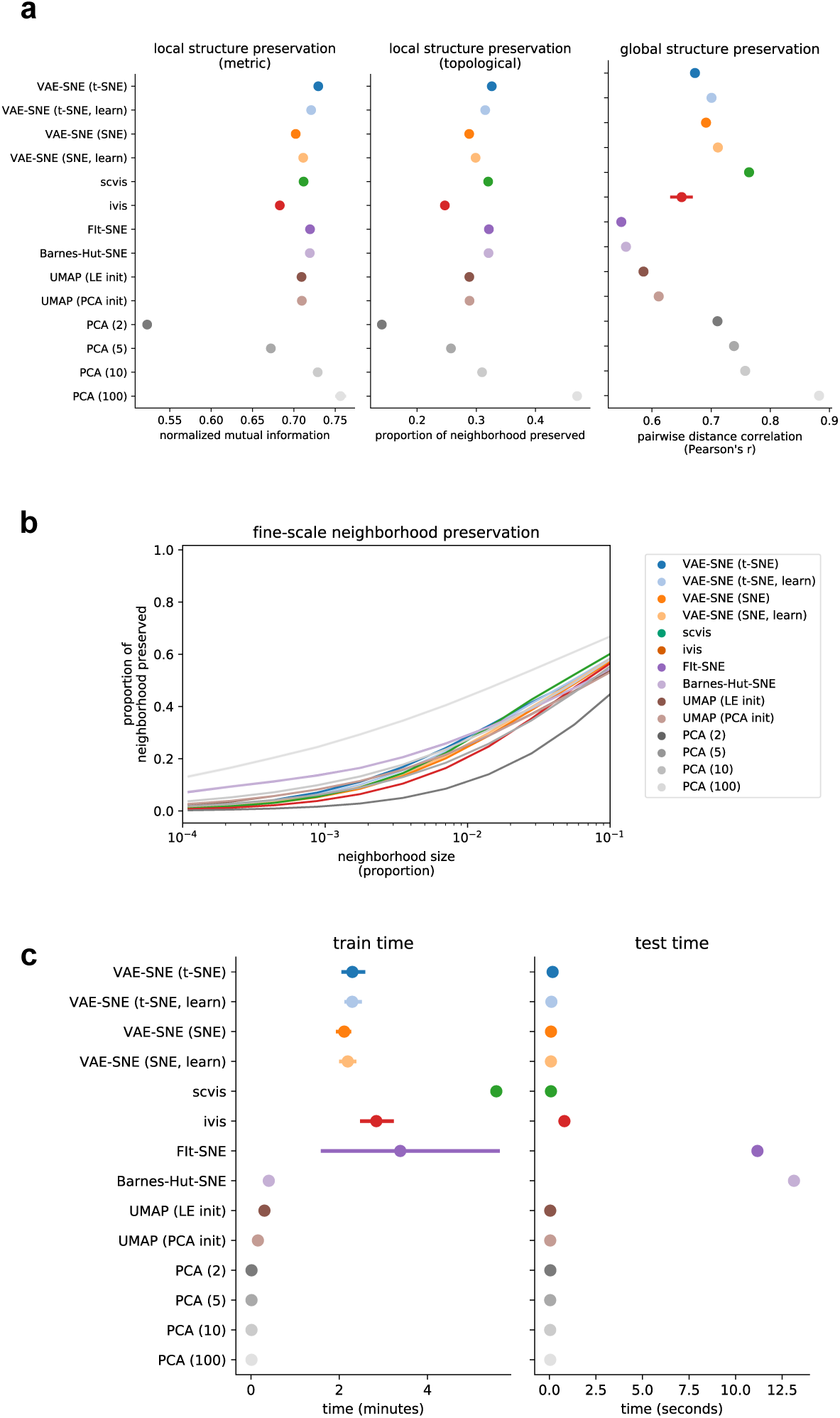
Dimension reduction performance for the single-cell RNA-seq dataset. Plots show performance comparisons for the single-cell RNA-seq dataset from La Manno et al. (2018) using the entire dataset. **a**, Mean and 95% interval of the bootstrap distribution for local and global structure preservation (for each metric n = 5 trials × 5 replicates = 25 per algorithm). **b**, Mean and 95% interval of the bootstrap distribution for fine-scale structure preservation across multiple neighbor sizes (as a proportion of the total embedding size; n = 14 neighborhood sizes × 5 trials × 5 replicates = 350 per algorithm). **c**, Elapsed time for embedding the training set and re-embedding the training set as a “test” set with each algorithm (for each metric n = 5 trials × 5 replicates = 25 per algorithm).

**Figure S6.**
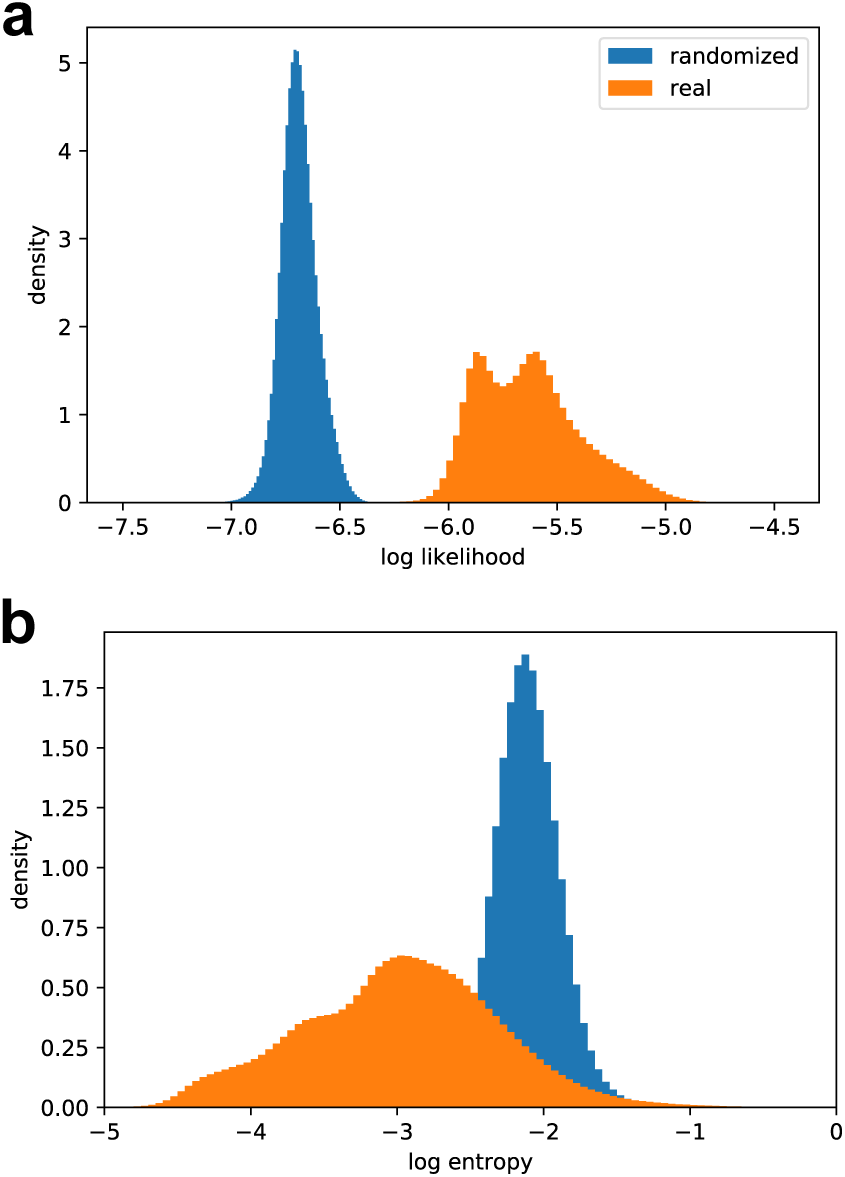
Likelihood and entropy distributions. **a**, Histograms of the log likelihood scores from the decoder (Eq. 1b; distortion) for real and randomized data (n = 232,000 for each distribution). **b**, Histograms of the log entropy from the approximate posterior (Eq. 12d) for real and randomized data (n = 232,000 for each distribution).

**Figure S7.**
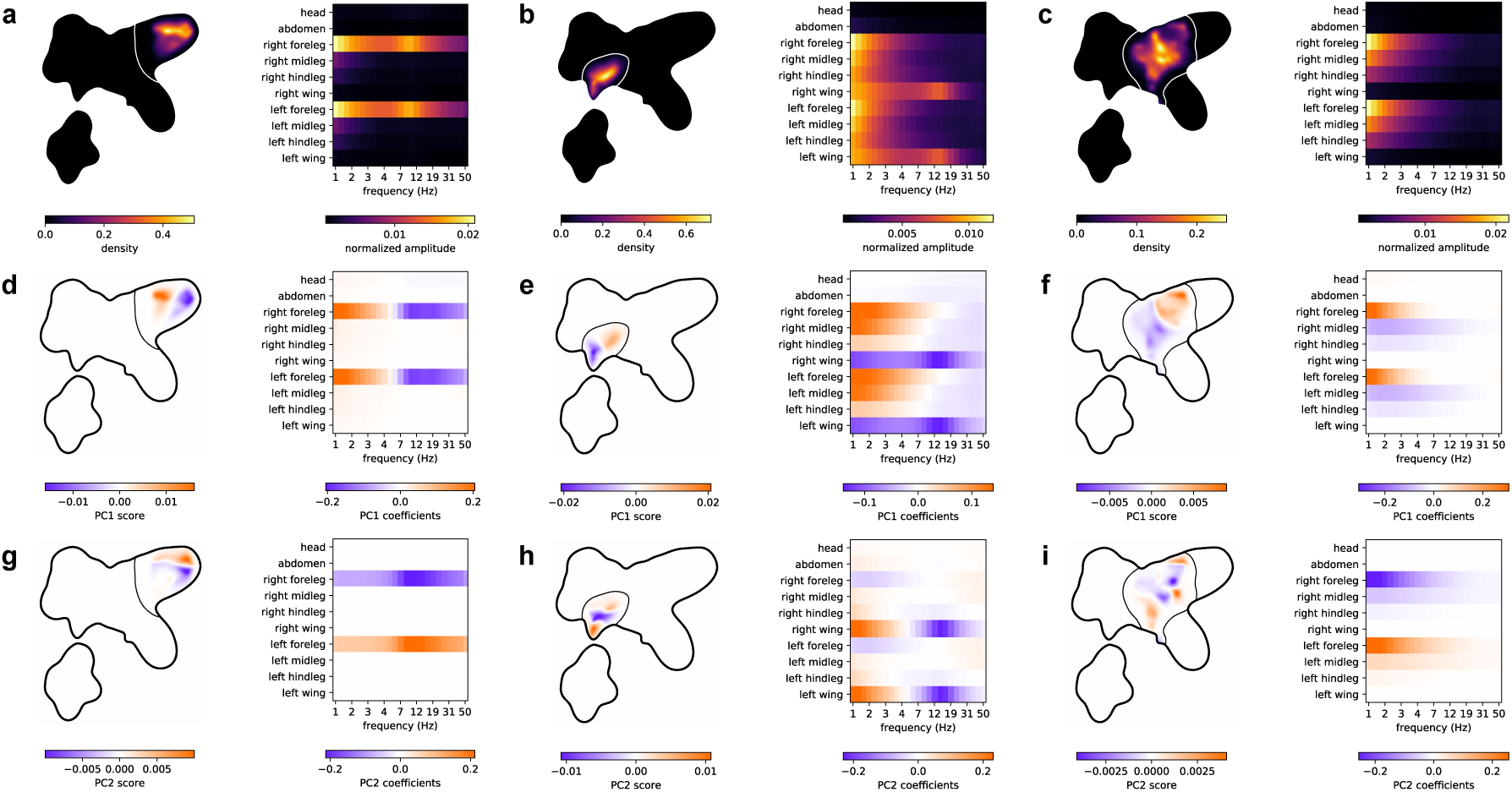
High-level behavioral clusters. Visualizations describing the manually-grouped high-level clusters for anterior grooming (**a**,**e**,**g**), wing movements (**b**,**e**,**h**) and small/slow leg movements (**c**,**f**,**i**). **a-c**, The 2-D posterior probability density for each cluster (left), where contours are the largest 90% probability density contour for each cluster distribution, and the mean spectrogram for each cluster (right). **d**-**i**, The principal component scores of the spectrograms assigned to each cluster visualized within the 2-D embedding (left) and the eigenvector coefficients describing the linear contribution of each spectrogram feature (right) for the principal component score.

**Figure S8.**
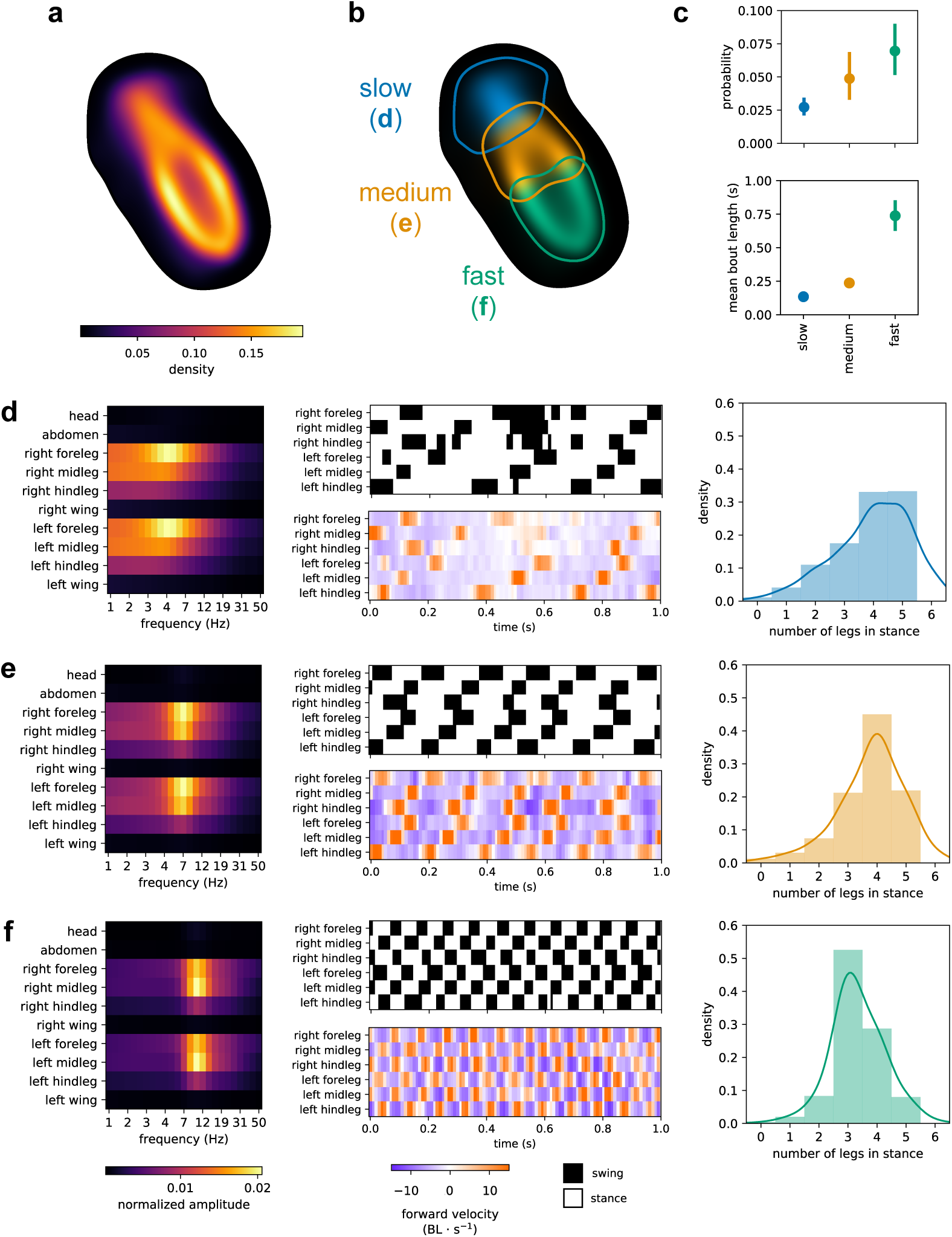
Low-level locomotion clusters. Visualizations describing the low-level clusters within the high-level locomotion cluster. **a-b**, The 2-D posterior probability density for the high-level cluster (**a**) and for each low-level cluster (**b**), where letters for each cluster label correspond to panels **d-f**. Contours are the largest 90% probability density contour for each cluster distribution. **c**, Mean and 95% bootstrap intervals of the marginal (stationary) probability and mean bout length for each low-level cluster (n = 59 per cluster). **d-f**, The mean spectrogram (left), example time segments (middle) showing forward velocity of each leg measured in body lengths (BL) per second and swing (forward velocity > 0 BL · s^−1^) or stance (forward velocity ≤ 0 BL · s^−1^) classification, and histograms (right) showing the number of legs classified as stance in each timestep assigned to each cluster (n = 0.57 million for slow, **d**; n = 1.03 million for medium, **e**; and n = 1.47 million for fast, **f**) — where the label for each panel in **d-f** corresponds to a cluster label in panel **b**. Example videos for these low-level clusters are shown in Video S2.

**Figure S9.**
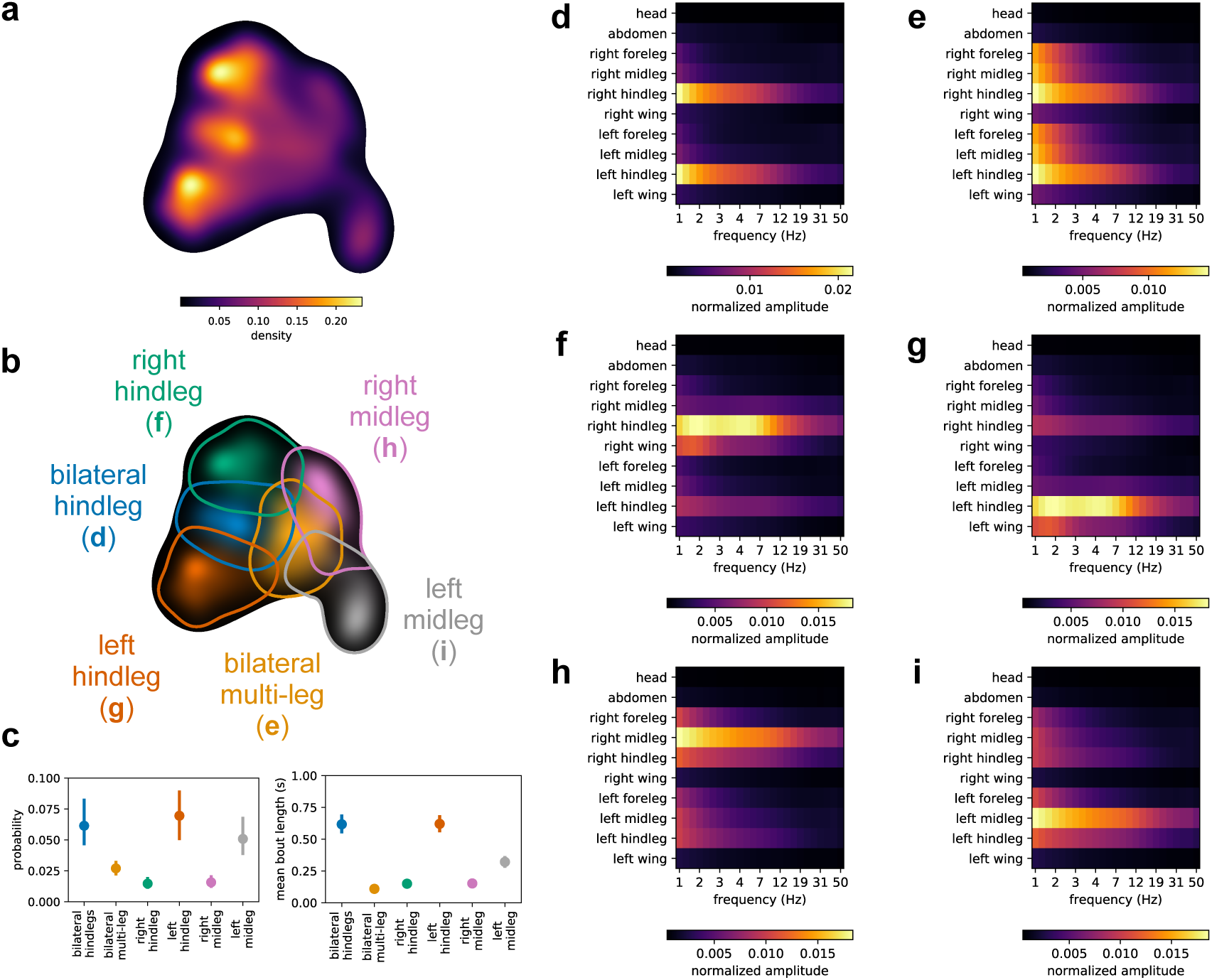
Low-level posterior grooming clusters. Visualizations describing the low-level clusters within the high-level posterior grooming cluster. **a-b**, The 2-D posterior probability density for the high-level cluster (**a**) and for each low-level cluster (**b**), where letters for each cluster label correspond to panels **d-i**. Contours are the largest 90% probability density contour for each cluster distribution. **c**, Mean and 95% bootstrap intervals of the marginal (stationary) probability and mean bout length for each low-level cluster (n = 59 per cluster). **d-i**, The mean spectrogram for each cluster — where the label for each panel in **d-i** corresponds to a cluster label in panel **b**. Example videos for these low-level clusters are shown in Video S4.

**Figure S10.**
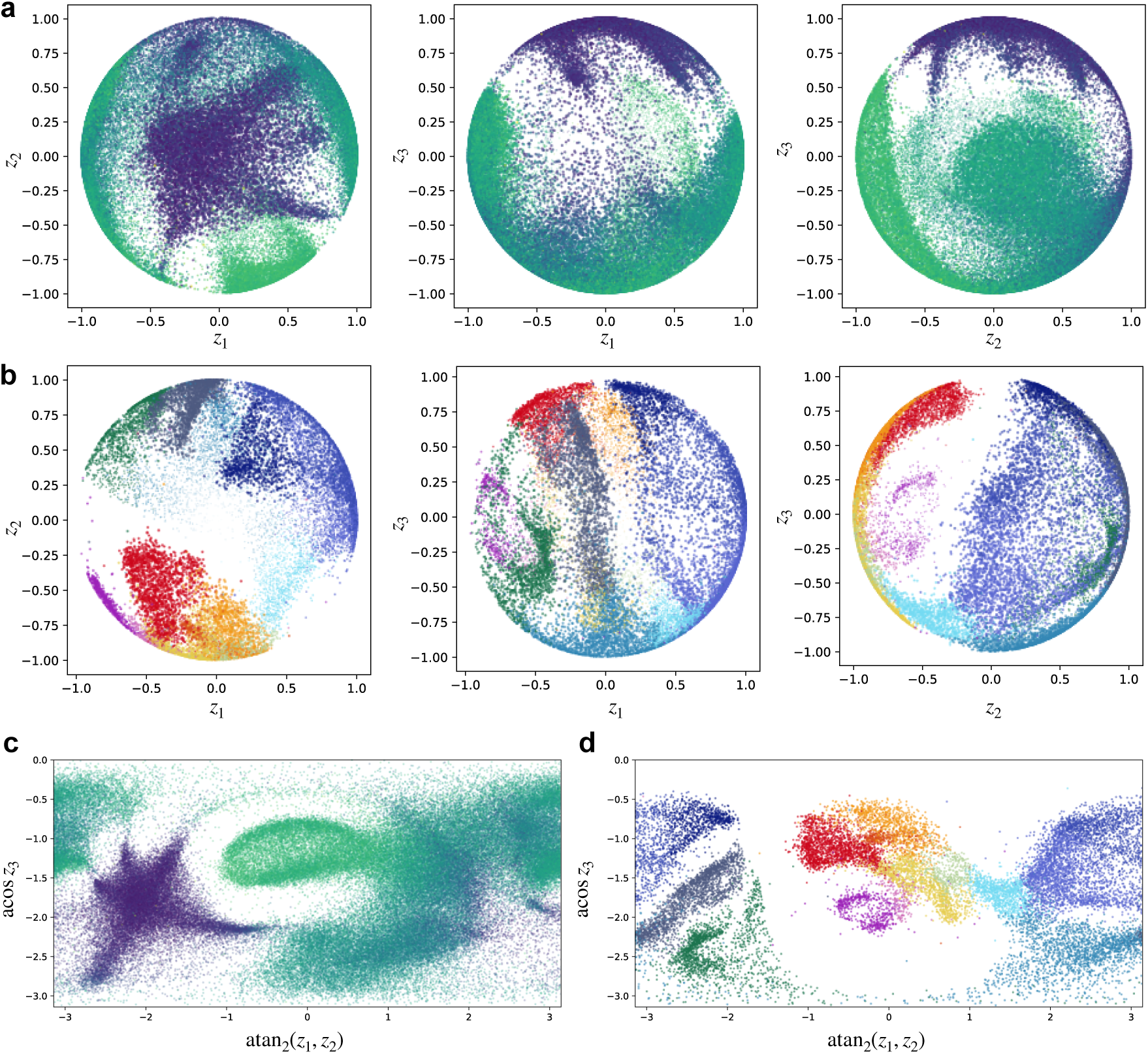
Spherical embeddings with von Mises-Fisher VAE-SNE. **a-b**, Spherical embeddings using VAE-SNE with a von Mises-Fisher similarity kernel (Appendix C.1) of the posture dynamics dataset (**a**; Video S8) from Berman et al. (2014, 2016); Pereira et al. (2019) and the single-cell RNA-seq dataset (**b**; Video S9) from La Manno et al. (2018). **c-d**, Stereographic (planar) projections of the spherical embeddings from **a-b**. Colors for **a-d** are the same as in Fig. 2 (total amplitude and cell type).

**Figure S11.**
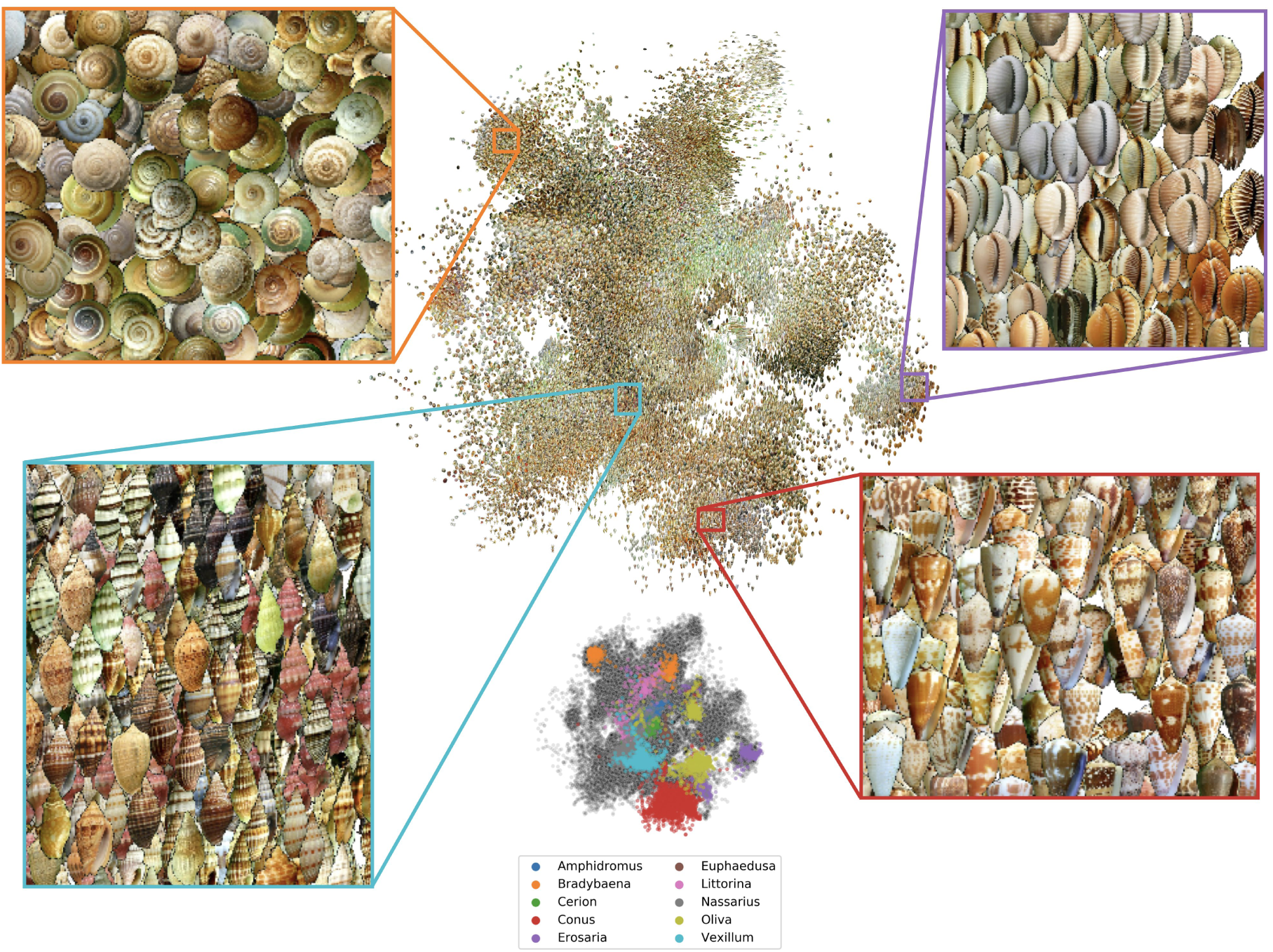
Embedding shell images. Shell images from Zhang et al. (2019) embedded in two dimensions using convolutional VAE-SNE. Insets illustrate example regions of perceptually similar images from the taxonomic genera *Bradybaena* (land snails; top-left), *Erosaria* (cowries; top-right), *Vexillum* (sea snails; bottom-left), and *Conus* (cone snails; bottom-right). Scatter plot (bottom-center) shows the 10 most common genera in the dataset.

**Figure S12.**
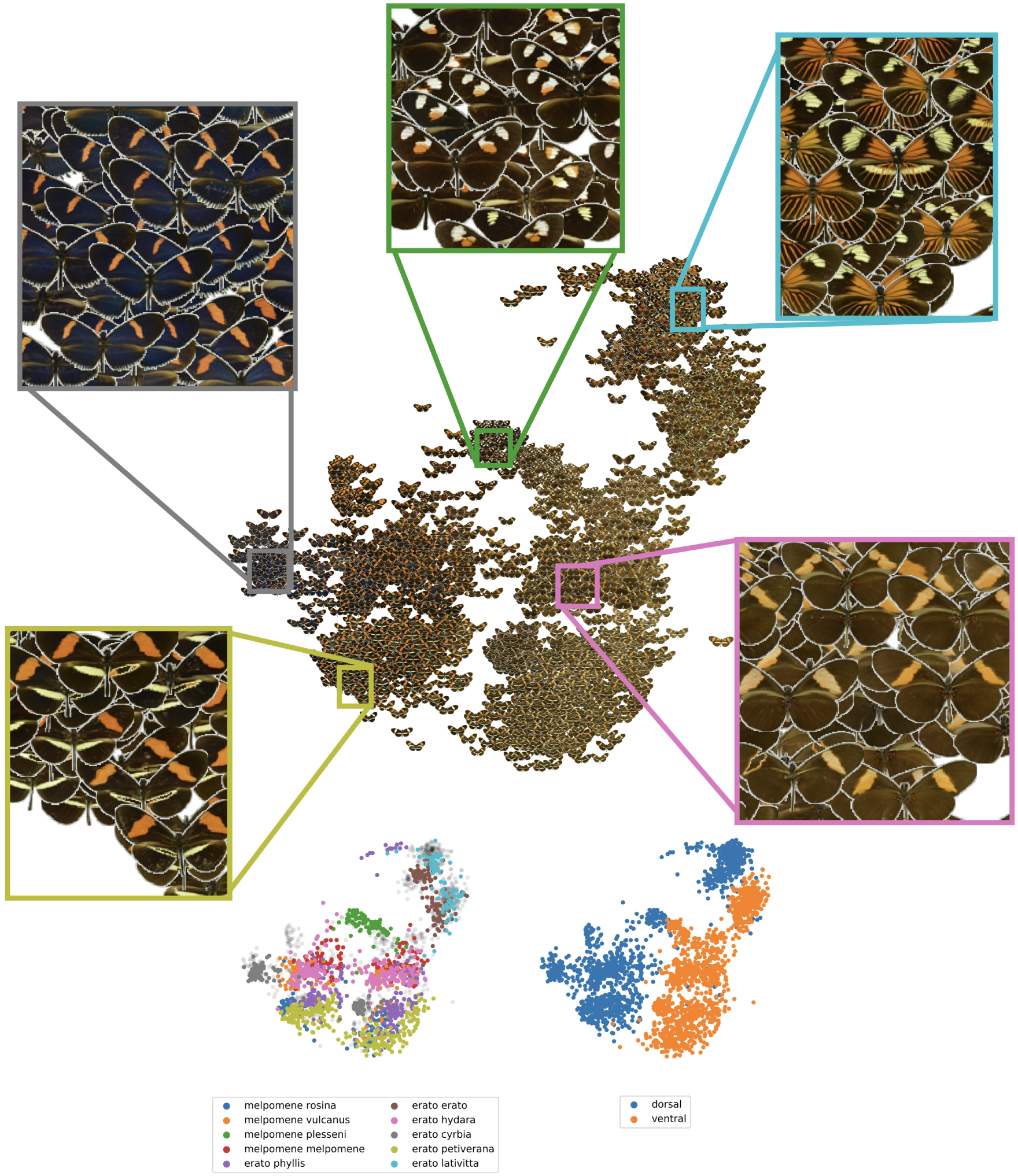
Embedding butterfly images. Butterfly (*Heliconius spp*.) images from Cuthill et al. (2019) embedded in two dimensions using convolutional VAE-SNE. Insets show example regions of perceptually similar subspecies (top). Scatter plots (bottom) show labels for the 10 most common subspecies in the dataset (bottom-left) and the image viewpoint relative to the specimen’s dorso-ventral body axis (bottom-right).

**Figure Video S1. Video segments labeled with VAE-SNE**. Randomly selected video segments (1*/*2 × speed) labeled with VAE-SNE illustrating the temporal dynamics of movements through the behavioral space and transitions between high-level clusters within the distribution. **a**, https://youtu.be/JlbSdKzvLfk; **b**, https://youtu.be/uWScG_UuzRQ; **c**, https://youtu.be/T8e_JSoCwMA

**Figure Video S2. Samples from the locomotion cluster**. Randomly sampled videos (1*/*3 × speed) from the locomotion cluster showing: **a**, slow walking (https://youtu.be/hB3JIRF2JGQ); **b**, medium walking (https://youtu.be/kNHGJypOGhs); and **c**, fast walking (https://youtu.be/A2sLtgYhHGc). Red lines show the posture tracking data for all 32 keypoints.

**Figure Video S3. Samples from the anterior grooming cluster**. Randomly sampled videos (1*/*3 × speed) from one of the anterior grooming clusters (https://youtu.be/0MT3lb2bJro). Red lines show the posture tracking data for all 32 keypoints.

**Figure Video S4. Samples from the posterior grooming cluster**. Randomly sampled videos (1*/*3 × speed) from the posterior grooming cluster showing: **a**, bilateral hindleg grooming (https://youtu.be/OTyf4pEQMo); **b**, right hindleg grooming (https://youtu.be/VTIwZp6d6b4); **b**, left midleg grooming (https://youtu.be/0vJvAINbfjw). Red lines show the posture tracking data for all 32 keypoints.

**Figure Video S5. Samples from the wing movements cluster**. Randomly sampled videos (1*/*3 × speed) from the wing movements cluster showing: **a**, wing extensions (https://youtu.be/lE31SeJ7ehY); and **b**, wing flicks (https://youtu.be/nsgnFbrk090). Red lines show the posture tracking data for all 32 keypoints.

**Figure Video S6. Samples from the small/slow leg movements cluster**. Randomly sampled videos (1*/*3 × speed) from the small/slow leg movement cluster showing: **a**, small leg movements (https://youtu.be/ARkH1uvPBnQ); **b**, slow leg movements (https://youtu.be/hwL7ovNjbBQ); **c**, small left midleg movements (https://youtu.be/o8vxtgwzx9Q) Red lines show the posture tracking data for all 32 keypoints.

**Figure Video S7. Samples from the idle cluster**. Randomly sampled videos (1*/*3 × speed) from the idle cluster (https://youtu.be/0wbdqmuCe_g). Red lines show the posture tracking data for all 32 keypoints.

**Figure Video S8. Spherical embedding of the posture dynamics dataset**. Rotating view of the posture dynamics dataset (https://youtu.be/QcDUlQUOvdo) from Berman et al. (2014, 2016); Pereira et al. (2019) embedded on a 3-D sphere using von Mises-Fisher VAE-SNE. Colors are the same as in Fig. 2 (total amplitude).

**Figure Video S9. Spherical embedding of the single-cell RNA-seq dataset**. Rotating view of the single-cell RNA-seq dataset (https://youtu.be/jyIWB6-qye0) from La Manno et al. (2018) embedded on a 3-D sphere using von Mises-Fisher VAE-SNE. Colors are the same as in Fig. 2 (cell type).

## A Variational autoencoders and the evidence lower bound

### A.1 VAEs as approximate Bayesian inference

As is common to most dimensionality reduction algorithms, we seek to model a high-dimensional data distribution *p*(**x**) using a low dimensional latent distribution *p*(**z**). Variational autoencoders (VAEs) are one such model that combines both modeling and inference by defining a joint distribution between a latent variable **z** and observed samples **x**. We can accomplish this using a generative model that maps samples from the low-dimensional latent distribution to the high-dimensional data distribution using a set of shared parameters ***θ***, which can take the form of a deep neural network model *p*_*θ*_(**x**|**z**) = DNN_*θ*_(**z**) with some prior over latent distribution *p*_*θ*_(**z**). We then wish to find the model parameters ***θ*** that maximize the joint likelihood, which can be written as:

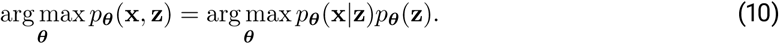

Although, to compute the low-dimensional distribution for the data, we then need to derive the latent posterior for the model *p*_*θ*_(**z**|**x**). This can be derived from the likelihood using Bayes’ rule:

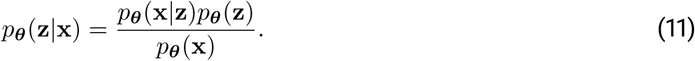

However, computing the integral in Eq. 11 *p*_*θ*_(**x**) = ∫ *p*_*θ*_(**x**|**z**)*p*_*θ*_(**z**) *d*z is not tractable in practice. Therefore, we require a way to approximate this latent posterior distribution, which is the exact problem for which VAEs provide a tractable solution.

Like other VAE models (Kingma and Welling, 2013; Kingma et al., 2014; Burda et al., 2015; Dilokthanakul et al., 2016; Ding et al., 2018; Dieng et al., 2019a), VAE-SNE performs dimensionality reduction by nonlinearly mapping observed high-dimensional data vectors **x** to a low-dimensional embedding **z** using a deep neural network (DNN) as an encoder function DNN_*ϕ*_ : **x** → **z** (Eq. 12e) with the goal of learning an approximate posterior over the latent distribution *q*_*ϕ*_(**z**|**x**) (Eq. 12d), where the parameters of the approximate posterior are learned as a function of the data (Eq. 12e) and the encoder parameters ***ϕ*** are then shared across observed samples — known as *amortization*. The model then maps latent vectors sampled from the low-dimensional embedding (Eq. 12c) to reconstruct the original high-dimensional space 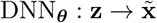 (Eq. 12a) using a generative decoder function we defined earlier (rewritten in Eq. 12b). More precisely:

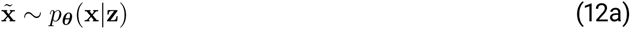

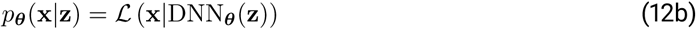

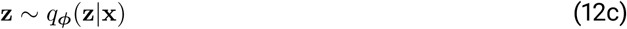

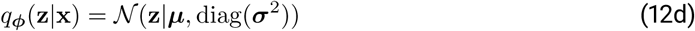

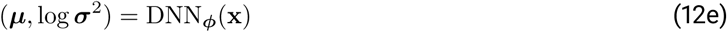

where ℒ(**x**|·) is a user-selected likelihood function parameterized by the decoder function DNN_*θ*_(**z**), and *N*(·|***µ***, diag(***σ***^2^)) is a multivariate Gaussian whose parameters ***µ*** and *σ*^2^ are a specified by the encoder function DNN_*ϕ*_(**x**).

### A.2 Deriving the evidence lower bound

After defining the generative model, we then wish to optimize the parameters of the encoder ***ϕ*** and decoder ***θ*** — given a set of observed samples from a data distribution **x** ∼ *p*(**x**) — so that the approximate posterior distribution *q*_*ϕ*_(**z**|**x**) matches closely with the true latent posterior from the generative decoder, or *q*_*ϕ*_(**z**|**x**) ≈ *p*_*θ*_(**z**|**x**). In other words, we wish to minimize the divergence between the two distributions, or:

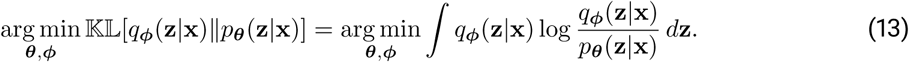

However, as we have already established, computing the true posterior is intractable, so researchers have derived a lower bound known as the evidence lower bound, or ELBO (Kingma and Welling, 2013), to approximate this objective. The ELBO can be derived directly from Eq.13(Adams, 2020), which is written as:

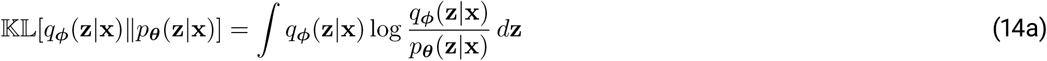

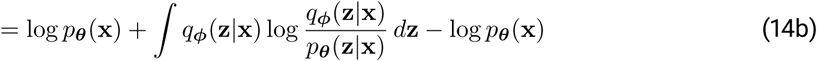

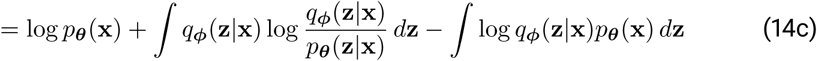

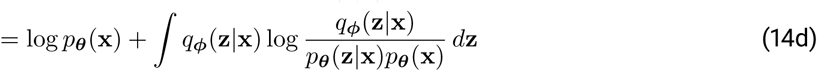

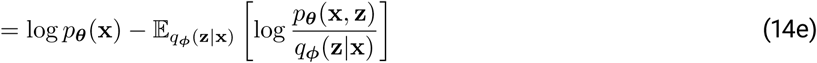

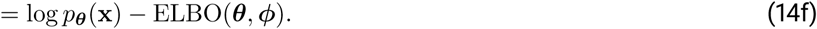

Because the Kullback-Leibler divergence is strictly non-negative, the ELBO is then a lower bound on the log marginal likelihood. However, The ELBO can also be derived by applying Jensen’s inequality, as is more common in the literature (Kingma and Welling, 2013), to directly calculate a lower bound on the log marginal likelihood, or:

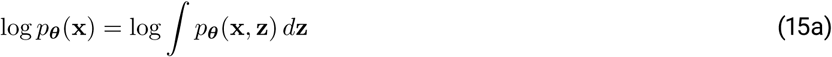

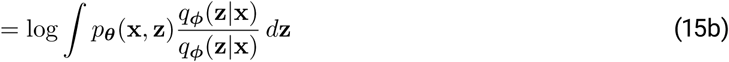

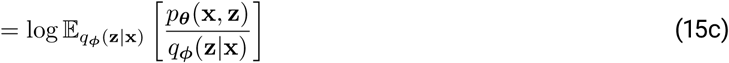

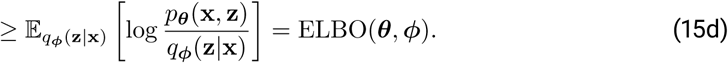

To learn the latent distribution given the model and the data, the ELBO is then maximized to optimize the model parameters. Here we write this as a minimization of the negative ELBO, which can be further decomposed into separate terms for the log-likelihood and the divergence between the approximate posterior and the prior over the latent distribution, or:

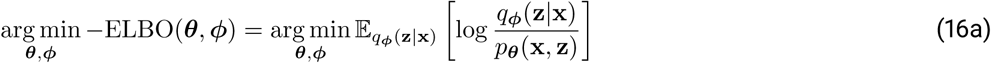

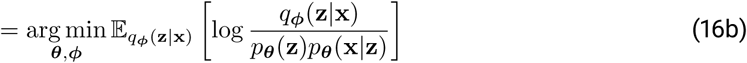

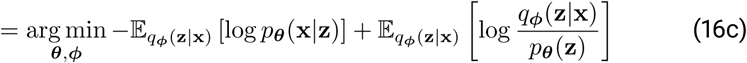

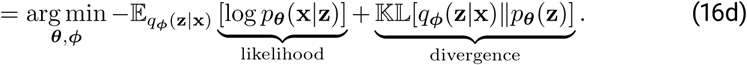

The derivation of the ELBO has also been discussed at length elsewhere (e.g., Kingma and Welling 2013; Kingma et al. 2014; Burda et al. 2015; Alemi et al. 2016; Dilokthanakul et al. 2016; Alemi et al. 2017; Ding et al. 2018; also see Kingma and Welling2019 for a comprehensive introduction).

### A.3 Importance-weighted ELBO

While we use only a single Monte Carlo sample from the approximate posterior per training batch, we also include a hyperparameter for multiple samples per training batch using the importance-weighted ELBO from Burda et al. (2015), which modifies how the expectation in Eq. 16c is calculated to produce a tighter bound on the loss by implicitly increasing the complexity of the posterior (Cremer et al., 2017). However, we did not see any obvious performance improvements when using the importance-weighted objective, and increasing the number of Monte Carlo samples per batch also increases training time. The general utility of calculating a tighter bound is also unclear (Rainforth et al., 2018) but this may be related to the generalization ability of the model. We leave further exploration of this hyperparameter for future work.

## B Stochastic neighbor regularization

For computing pairwise similarities, we largely follow Hinton and Roweis(2003) and van der Maaten and Hinton(2008) by modeling local neighborhoods as the probability of transitioning from a landmark point to its nearby neighbors when performing a random walk initialized from the landmark. By modeling local neighborhoods as probability distributions and then minimizing the divergence between the neighborhood distributions in high- and low-dimensional space, we preserve more local structure within the low-dimensional embedding than a standard VAE (Ding et al., 2018).

### High-dimensional transition probabilities

To accomplish this, pairwise transition probabilities in high-dimensional space *t*(**x**_*j*_|**x**_*i*_) are modelled by applying a Gaussian kernel to convert the pairwise distances between data points *d*(**x**_*i*_, **x**_*j*_) into conditional probabilities — with self transitions set to *t*(**x**_*i*_|**x**_*i*_) = 0. While Ding et al. (2018) use these asymmetric conditional probabilities *t*(**x**_*j*_|**x**_*i*_) directly for the high-dimensional similarities, van der Maaten and Hinton (2008) show that symmetrizing the pairwise similarities so that *p*(**x**_*j*_|**x**_*i*_) = *p*(**x**_*i*_|**x**_*j*_) reduces susceptibility to outliers, which can become ill-determined in the low-dimensional embedding with an asymmetric kernel. Therefore, we use the symmetrized conditional probabilities, which are computed as:

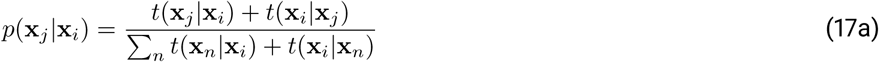

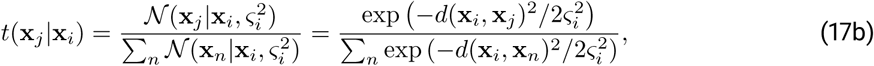

for *n* = 1, …, *N* and *n* ≠ *i*, where *d*(·, ·) is a user-selected distance metric, such as the Euclidean distance. The landmark data point **x**_*i*_ can then be considered the mean, and 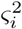 is the variance of the Gaussian kernel describing the local neighborhood around **x**_*i*_ — thereby assigning more probability mass to nearby neighbors. The variance 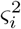 is selected for each data point via binary search such that 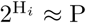, where P is the desired perplexity (a user-defined hyperparameter), 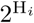 is the perplexity of the kernel for the *i*th data point, which approximately corresponds to the number of nearest neighbors considered by the kernel, and H_*i*_ is the Shannon entropy in bits, or:

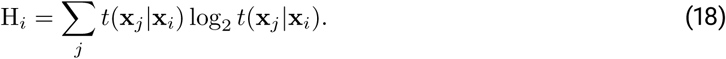

### Low-dimensional transition probabilities

The low-dimensional similarities *q*_*ϕ*_(**z**_*j*_ **z**_*i*_) are then calculated according to Hinton and Roweis (2003) and van der Maaten and Hinton (2008) using a kernel function *w*_*ϕ*_(**z**_*j*_|**z**_*i*_) to convert pairwise distances into conditional probabilities:

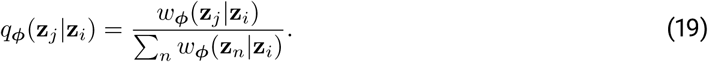

As in high-dimensional space, self transitions are set to *q*_*ϕ*_(**z**_*i*_|**z**_*i*_) = 0. Here we test two kernel functions for preserving Euclidean similarities.

### t-SNE kernel

First is the heavy-tailed Student’s *t*-distributed kernel used for the t-SNE algorithm (van der Maaten and Hinton, 2008) with the log probability function written as:

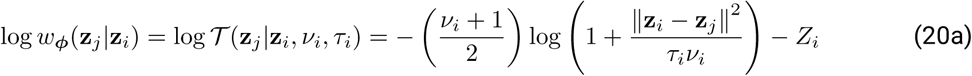

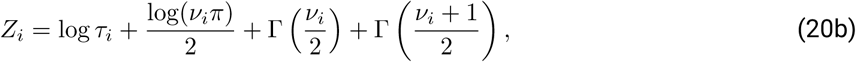

where *τ*_*i*_ is the scale, *ν*_*i*_ is the degrees of freedom, which varies the heavy-tails of the kernel, and Γ(·) is the gamma function. We write this as a log probability to more clearly show the relationship with the similarity loss term derived later in this section (Eq. 23c). The Student’s *t*-distribution is used primarily to alleviate the “crowding problem” (van der Maaten and Hinton, 2008) that can occur with other nonlinear embedding algorithms, including the original SNE algorithm (Hinton and Roweis, 2003), where points are too densely packed in the low-dimensional space and moderately distant points are “crushed” together as an artifact of the embedding algorithm.

### SNE kernel

Secondly, we test a Gaussian kernel — the kernel used for the original SNE algorithm (Hinton and Roweis, 2003; van der Maaten and Hinton, 2008) — with the log probability function:

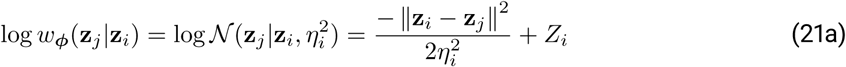

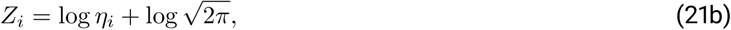

where 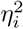 is the variance.

### Setting the kernel parameters

The kernel parameters for the low-dimensional similarities are typically set to a constant value, such as *τ*_*i*_ = *ν*_*i*_ = *η*_*i*_ = 1 (van der Maaten and Hinton, 2008), or are scaled linearly with the dimensionality of the latent embedding (van der Maaten, 2009), but we also test similarity kernels where these parameters are learned for each data point, parameterized by the encoder DNN_*ϕ*_(**x**) — an idea proposed by van der Maaten(2009). When the kernel parameters are constant across all data points, the log normalization terms (Eqs. 20b, 21b) used for calculating the log probabilities can be omitted as an additive constant that has no effect on the calculations after normalization. However, this term is potentially important for optimization when learning these parameters as a function of each data point, so we include it in our calculations.

### Reinterpreting the similarity loss term

To maximize numerical stability when optimizing the similarity term, we substitute the cross-entropy between the high-dimensional and low-dimensional similarities H[*p*(**x**_*j*_|**x**_*i*_), *q*_*ϕ*_(**z**_*j*_|**z**_*i*_)], which is proportional to the Kullback-Leibler divergence and, after dropping the expectation, can be derived as follows:

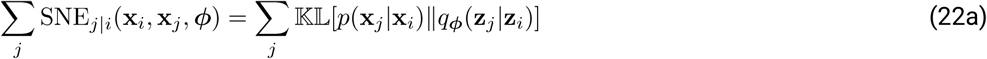

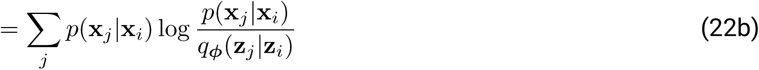

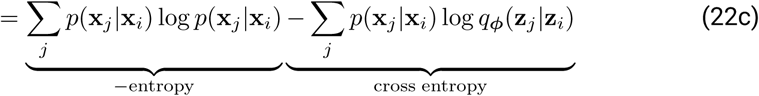

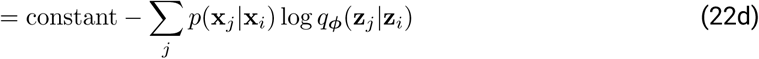

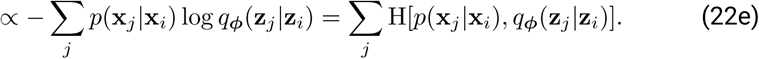

Consequently, the Kullback-Leibler divergence for the similarity term can be reinterpreted as the cross-entropy between the pairwise similarities up to an additive constant (the negative entropy of the high-dimensional similarities), which can be omitted for the purposes of optimization. To further improve numerical stability for this computation, the cross-entropy is decomposed into attractive and repulsive forces using the unnormalized similarities (following Ding et al. 2018; Kobak and Berens 2019), which is written as:

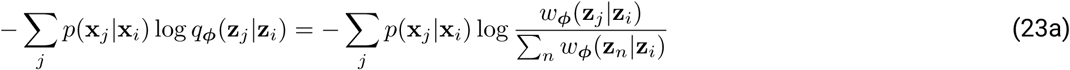

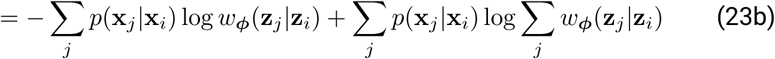

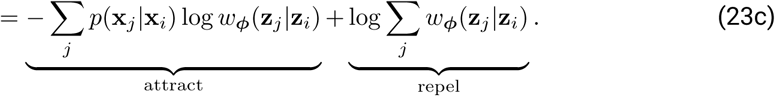

This may also help to clarify why we wrote the low-dimensional kernels as log-probability functions in Eqs. 20a, 21a.

## C Extensions of VAE-SNE

### C.1 Spherical embeddings with a von Mises-Fisher kernel

In addition to embeddings with Euclidean geometry, we introduce a version of VAE-SNE that uses polar geometry and embeds high-dimensional data on the surface of a 3D unit sphere. We calculate the high-dimensional similarities according to Appendix B, but we alter the calculations for the transition probabilities by using the cosine similarity for the high-dimensional pairwise metric. After normalization, this is equivalent to using a (hyper)spherical von Mises-Fisher distribution as the similarity kernel, or:

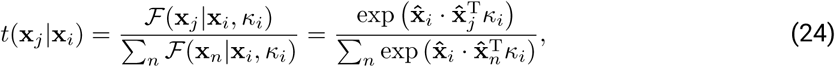

where 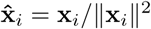 and *κ*_*i*_ is the concentration parameter (the inverse variance 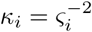), which is selected using binary search to match the perplexity to a desired value (see Appendix B for details). We then calculate the low-dimensional similarities using a 3D von Mises-Fisher kernel to create a spherical embedding:

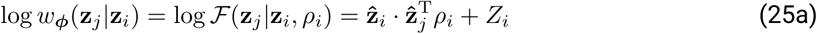

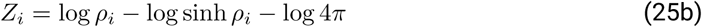

where 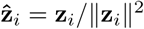 and *ρ*_*i*_ is the concentration parameter (inverse variance). The log normalization term (Eq. 25b) can be omitted when *ρ*_*i*_ is set to a constant, but we include it for the purposes of optimizing *ρ*_*i*_ as a function of each data point.

The idea of using spherical embeddings for dimensionality reduction has been explored previously with the von Mises-Fisher stochastic neighbor embedding (VMF-SNE) algorithm (Wang and Wang, 2016) as well as more recent work by Ding and Regev (2019) who apply this type of embedding to visualize single-cell RNA-seq data. The UMAP algorithm (McInnes et al., 2018) has a similar option to embed data in polar coordinates, as well as other non-Euclidean spaces. VAEs with (hyper)spherical latent variables have also been explored extensively in the machine learning literature (Davidson et al. 2018; reviewed by Ding and Regev2019). This type of spherical representation can be useful for data analysis, as high-dimensional vectors are often more accurately represented in polar coordinates. Similar to a heavy-tailed Student’s *t* similarity kernel (van der Maaten and Hinton, 2008), a spherical von Mises-Fisher similarity kernel can also prevent “crowding” of the data toward the center of the latent coordinate system (Davidson et al., 2018; Ding and Regev, 2019), which is undesirable for visualizing data (van der Maaten and Hinton, 2008). To test this extension, we use von Mises-Fisher VAE-SNE to embed the posture dynamics dataset from Berman et al. (2014, 2016); Pereira et al. (2019) as well as the single-cell RNA-seq dataset from La Manno et al. (2018) and visualize the embeddings across the three dimensions of the unit sphere (Fig. S10; Video S8;Video S9). We find that the results are qualitatively similar to 2-D Euclidean embeddings of the same data (Fig. 2), but are instead embedded across a 3-D sphere. Despite not using a heavy-tailed similarity kernel (van der Maaten and Hinton, 2008) these spherical embeddings naturally do not exhibit any crowding problems (Davidson et al., 2018; Ding and Regev, 2019), which may make this a useful visualization tool for some scenarios.

### C.2 Convolutional VAE-SNE for image data

We introduce a convolutional version of VAE-SNE for embedding image data from raw pixels. This version of VAE-SNE is modified by first applying a 2-D convolutional neural network CNN_*ϕ*_ — a SqueezeNet v1.1 (Iandola et al., 2016) pretrained on ImageNet (Deng et al., 2009) — to each image and then calculating the pairwise similarity using spatially-pooled feature maps from the CNN_*ϕ*_ output. The high-dimensional transition probabilities (Appendix B) are then calculated using a Gaussian kernel:

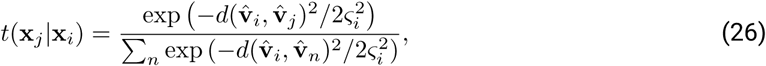

where 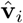 is a vector of spatially-pooled feature maps from the CNN_*ϕ*_ output, or 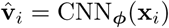. The approximate posterior is then calculated as a nonlinear function of the pooled feature maps 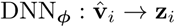, which is written as 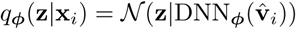. For the decoder we use a feed-forward network 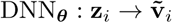 as before, where 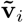 is a reconstruction of the CNN_*ϕ*_ output 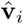. We then apply mean squared error between the pooled feature maps and the reconstruction as the likelihood function for the distortion loss (Eq. 1b). A convolutional decoder could also be used to fully reconstruct the raw image pixels, but we found simply reconstructing the pooled feature maps to be effective for visualizing the distribution of images in two dimensions.

To demonstrate the utility of convolutional VAE-SNE, we embed natural history image datasets of both shells (Zhang et al., 2019) and (*Heliconius spp*.) butterflies (Cuthill et al., 2019). We then visualize these embeddings to qualitatively assess performance of this VAE-SNE variant (Figs. S11, S12). We find that perceptually similar images are grouped together in the embedding based on complex sets of image features — rather than simple heuristics like color — and these groupings correspond to taxonomic relationships within the dataset, which were not explicitly included as part of the training set. This variant of VAE-SNE is functionally similar to using the perceptual distance (Johnson et al., 2016a; Wham et al., 2019) as a similarity metric and likelihood function except that the model can be trained end-to-end with small batches of images directly using raw pixels instead of first preprocessing images to produce feature activations. These results demonstrate that VAE-SNE can be used to analyze very large image datasets, by loading images in small batches, and can also be extended to images with variable resolution, by integrating across feature map outputs from the CNN to remove the spatial dimension — both of which are typically not possible with other dimension reduction algorithms.

